# Parasitic worm-derived ES-62 promotes health- and life-span in high calorie diet-fed mice

**DOI:** 10.1101/622753

**Authors:** Jenny Crowe, Felicity E. Lumb, James Doonan, Margaux Broussard, Anuradha Tarafdar, Miguel A. Pineda, Carmen Landabaso, Lorna Mulvey, Paul A. Hoskisson, Simon A. Babayan, Colin Selman, William Harnett, Margaret M. Harnett

**Affiliations:** Institute of Infection, Immunity and Inflammation, University of Glasgow, Glasgow G12 8TA, UK; Strathclyde Institute of Pharmacy and Biomedical Sciences, University of Strathclyde, Glasgow, G4 0RE, UK; Glasgow Ageing Research Network (GARNER), Institute of Biodiversity, Animal Health and Comparative Medicine, University of Glasgow, Glasgow, G12 8QQ, UK

**Author notes:** Joint first authors (Jenny Crowe, Felicity E. Lumb). Corresponding authors: Margaret Harnett, Institute of Infection, Immunity and Inflammation, College of Medical, Veterinary and Life Sciences, University of Glasgow, Glasgow G12 8TA, UK; e.mail –; William Harnett, Strathclyde Institute of Pharmacy and Biomedical Sciences, 161 Cathedral Street, University of Strathclyde, Glasgow G4 0RE, UK; e.mail.

## Abstract

The recent extension of human lifespan has not been matched by equivalent improvements in late-life health due to the global pandemic in type-2 diabetes, obesity and cardiovascular disease, ageing-associated conditions exacerbated by widespread adoption of the high calorie Western diet (HCD). As a novel therapeutic strategy, we have investigated the potential of ES-62, an immunomodulator secreted by the parasitic worm *Acanthocheilonema viteae*, to improve healthspan by targeting the chronic inflammation that drives metabolic dysregulation underpinning ageing-induced ill-health. We found that ES-62 improves a range of inflammatory but also pathophysiological, metabolic and microbiome parameters, when administered subcutaneously at only 1 µg/week throughout the lifespan of HCD-fed mice. Strikingly, ES-62 induced sex-specific healthspan signatures and indeed, it substantially increased the median survival of male, but not female, HCD-mice. Modelling of 113 responses contributing to these differential signatures by machine learning approaches now signposts candidate parameters key to promoting both healthspan and lifespan.

---

Adoption of the modern Western diet (high fat and high sugar) has significantly contributed to the global major public health problems of metabolic syndrome, obesity, type-2 diabetes and cardiovascular disease; ageing-associated comorbidities that impact on both wellbeing (healthspan) and lifespan^1, 2^. Interestingly, epidemiological data suggest that such high calorie diet (HCD)-associated diseases are rising fastest in in the developing world^2, 3^, regions where parasitic worms (helminths) and other infectious agents are being eradicated^4^. Helminths promote their survival by releasing excretory-secretory (ES) products that, by dampening inflammation and promoting tissue repair, act to prevent worm expulsion but also limit host pathology^4, 5^. Thus, the relatively rapid eradication of these parasites appears to have resulted in hyper-active host immune systems, characterised by chronic low-grade inflammation that may (further) contribute to development of obesity and associated metabolic syndrome co-morbidities as well as their reciprocal risk factors, allergic and autoimmune inflammatory disorders^6^, in developing and urbanised countries^4, 5^. Whilst this has questioned the wisdom of current mass parasite eradication programs, it has emphasised the potential of utilising worm infections or ES to treat a wide range of non-communicable diseases that are characterised by chronic inflammation^7–9^.

We have previously shown that a single, defined ES protein, ES-62, can resolve chronic inflammation by normalising aberrant MyD88 signalling to homeostatically restore immunoregulation, irrespective of the inflammatory phenotype^4, 5, 10–12^. MyD88 is increasingly recognized as a key innate receptor transducer and integrator of dysregulated inflammatory and metabolic pathways^7, 13–15^. Indeed, adoption of a HCD induces reprogramming of innate immune responses, with the resulting chronic TLR4/MyD88 signalling^16^ playing critical roles in promoting generation of pro-inflammatory (M1-like CD11c^+^) macrophages, glucose intolerance, β-cell failure and consequent inflammation of metabolic (adipose, pancreas and liver) tissues^16, 17^. Critically, these HCD-TLR4/MyD88-induced effects are exacerbated by ageing^18^. Thus, based on ES-62’s targeting of MyD88 and demonstrable ability to protect against chronic inflammatory allergic and autoimmune conditions^4, 5, 10–12^, we investigated whether it could safeguard against the impact of HCD-associated ageing in C57BL/6J mice. Our approach of combining longitudinal (survival) and cross-sectional (intervention) studies has revealed that ES-62 differentially improves healthspan in both male and female HCD-fed mice and significantly extends median lifespan of male HCD-fed mice. Of additional interest, this strategy has identified dynamic and differential diet-, sex- and age-associated immunological and metabolic healthspan signatures.

## Results

### ES-62 extends lifespan in male C57BL/6J mice fed a HCD

ES-62 did not significantly increase the median lifespan of a mixed sex cohort of HCD-fed C57BL/6J mice (Figure 1a). However, and consistent with previous findings that interventions that impact on health- and lifespan often exhibit sexual dimorphism^19–21^, Cox regression analysis revealed significant sex differences. Thus, ES-62 substantially extended the median lifespan of male (PBS, 629; ES-62, 703 days), but not female (PBS, 653; ES-62, 629 days) HCD-mice (Figure 1b-e; Supplementary Table 1).

**Figure 1:**
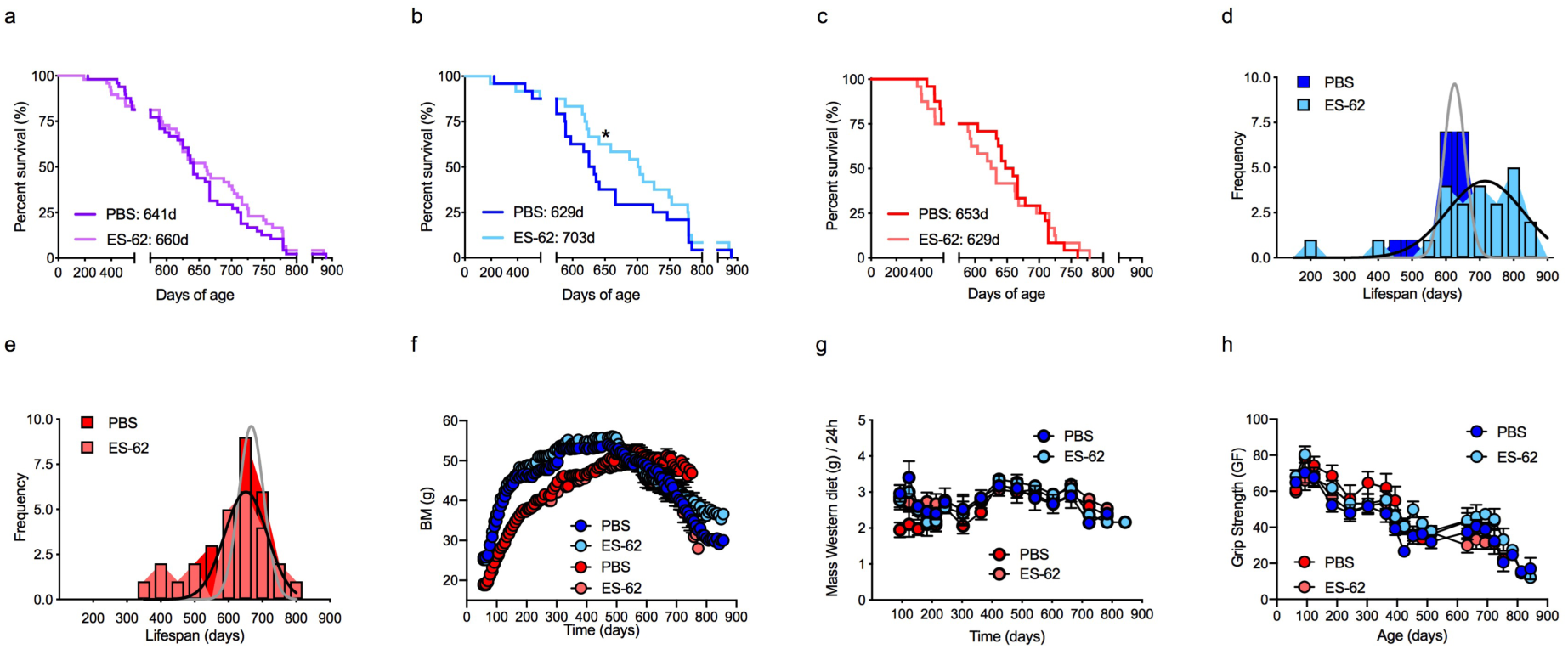
ES-62 extends lifespan in male HCD-fed C57BL/6J mice. Kaplan-Meier survival curves (a-c) for HCD-fed mice treated either with PBS or ES-62 were analysed as (a) mixed sex cohorts (PBS, n=48; ES-62, n=48); (b) male mice (Blue lines; PBS, n=24; ES-62, n=24, *p<0.05) only and (c) female mice (Red lines; PBS, n=24; ES-62, n=24) only. Median survival values for the relevant groups are shown as figure inserts. The data for the male and female cohorts are also presented as the frequency distribution of lifespan (d, male; e, female). Longitudinal analysis of body mass (BM; f), food intake (g) and grip strength (h) measurements was undertaken where data are presented as mean values ± SEM of the individual mice (Male, blue symbols; Female, red symbols) surviving at the indicated time-points.

Low body mass (BM), particularly in early life, is associated with longevity in genetically heterogeneous mice, fed a normal chow diet^22^. Notably therefore, the extension of life-span in male ES-62-HCD-fed mice did not reflect prevention of HCD-increased BM as ES-62 did not significantly reduce either peak BM (Male: PBS, 56.20 ± 0.93; ES-62, 57.22 ± 1.10 g; Female: PBS, 53.19 ± 1.32, ES-62, 50.77 ± 1.17 g) or BM over the course of weight gain (Figure 1f). Of note, although BM declined sharply after reaching maximal levels in male HCD (PBS and ES-62)-mice, female HCD-PBS mice maintained a relatively stable body weight until death: however, ES-62-treated female HCD mice exhibited a similar decline in weight to that of the aged male HCD cohorts (Figure 1f). This did not reflect changes in food intake, which was not altered by ES-62 in either male or female HCD-fed mice (Figure 1g).

Mirroring findings that maximum BM is not a good predictor of longevity in mice fed a normal diet^22^, there was no correlation between the peak BM achieved by individual mice and their longevity in any of the cohorts (Supplementary Figure 1a & b). Moreover, the significant association (peaking about 150 days of age) between low BM and longevity reported in chow-fed mice when measured between 60 and 720 days of age^22^, was not recapitulated in any of the HCD-fed cohorts when BM was measured at 116, 160, 340 or 500 days of age (Supplementary Figure 1c-j). Rather, perhaps critically, ES-62 appeared to protect against the loss of weight generally occurring during late ageing (>720-day old mice) in male, but not female, HCD-PBS mice (Figure 1f). As typically, lean mass declines whilst fat mass increases with age, this may suggest that ES-62 acts to preserve lean mass and prevent sarcopenia: perhaps supporting this, the negative correlation between BM at death and longevity in male, but not female, HCD-PBS mice is abolished by treatment with ES-62 (Supplementary Figure 1k & l). However, there was no corresponding protection afforded by ES-62 against decline in grip strength, one of the indicators of age-associated frailty (Figure 1h).

As expected given their life-long exposure to an energy-rich diet, many of the mice within our lifespan study exhibited evidence of liver tumours upon post mortem irrespective of sex or treatment (Supplementary Table 2). We therefore addressed the impact of ES-62 on metabolic tissues and their associated dysregulated functional responses in a series of cross-sectional studies (at d160, d340 and d500) comparing male and female HCD-fed mice with young (day 56) and aged-matched normal chow-fed control cohorts.

### ES-62 and the impact of HCD on BM and metabolic tissues

Mice in the cross-sectional cohorts exhibited very similar kinetics of BM gain to those in the longevity study, with analysis of their mean BM values at cull independently confirming that ES-62 had no significant effect on HCD-induced obesity (Supplementary Figure 2a-j). Analysis of metabolic organ mass (normalised to total BM; Supplementary Figure 2k-p) revealed that HCD-fed mice exhibit substantial increases in their levels of gonadal and retroperitoneal visceral fat during ageing, relative to their normal chow controls. However, consistent with the loss of metabolic regulation occurring during ageing^23^, mice fed a normal diet also showed pronounced increases in these fat depots, particularly in the d500 cohorts (Supplementary Figure 2k-n). Exposure to ES-62 generally had marginal effects on metabolic organ size in HCD-fed mice. However, the livers from the male HCD-ES-62 cohort were found to be significantly smaller than those from their (PBS) control group at d500 although this sex effect may reflect that the HCD had little overall effect on liver mass in obese relative to chow-fed female mice (Supplementary Figure 2o & p).

### ES-62 protects adipocyte health in HCD-fed mice

Deeper analysis revealed that, as expected, male and female mice fed a HCD exhibited pronounced adipocyte hypertrophy (Figure 2a-h), but not hyperplasia (Supplementary Figure 3a-d), in both gonadal and retroperitoneal visceral fat deposits, relative to their normal chow controls and this was evident by the day 160 timepoint. Adipocytes in gonadal fat from either the male or female chow-fed cohorts showed only a slow progressive increase in size with age whereas those in retroperitoneal fat were not significantly different from their HCD-PBS counterparts at d500, with clear increases in adipocyte size being detected by d160 in the male chow-fed cohort (Figure 2a-h). Interestingly therefore, whilst ES-62 was able to reduce the hypertrophy observed in both gonadal and retroperitoneal visceral fat depots in male HCD mice, it was only able to delay the HCD-increase in adipocyte size in retroperitoneal fat and had no effect on gonadal adipocyte area in female HCD-mice. Indeed, in male mice, the retroperitoneal, but not gonadal, adipocytes in ES-62-treated HCD-fed mice more closely resembled the morphology of those from the chow-fed rather than HCD-PBS cohorts (Figure 2a-h). Collectively, these data suggest that ES-62 may act to protect adipocyte metabolic function, particularly in male HCD-mice.

**Figure 2.**
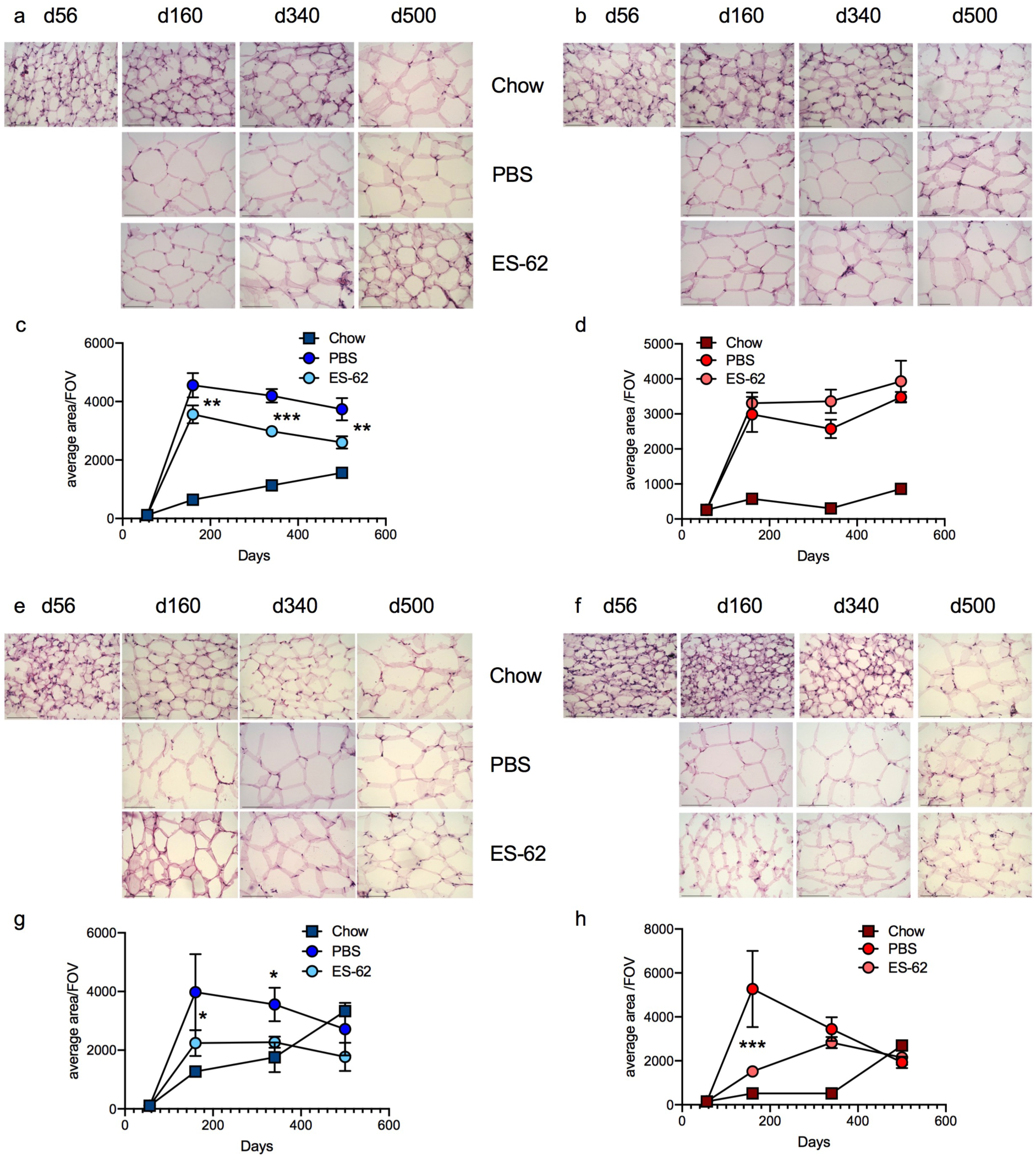
ES-62 ameliorates visceral adipocyte hypertrophy in HCD-fed mice. Representative images (scale bar 100 µm) of gonadal fat from male (a) and female (b) chow- and HCD- (PBS or ES-62-treated) mice stained with H & E and resultant quantitative analysis of adipocyte size where data are presented as the mean values ± SEM, where n=4-5 individual male (c) and female (d) mice at each time point and the values for each mouse are means derived from n=3 replicate analyses. Representative images (scale bar 100 µm) of retroperitoneal fat from male (e) and female (f) chow- and HCD-(PBS-or ES-62-treated) mice stained with H & E and resultant quantitative analysis of adipocyte size where data are presented as the mean values ± SEM, where n ≥ 4 individual male (g) and female (h) mice and the values for each mouse are derived from n=3 replicate analyses. For clarity, only significant differences between the HCD-PBS and HCD-ES-62 cohorts are shown on the figures, where significance is denoted by *p < 0.05, **p < 0.01 and ***p < 0.001.

To investigate the potential mechanisms involved, we examined the effect of ES-62 on eosinophil levels in the gonadal and retroperitoneal visceral fat depots. Eosinophils have been proposed to maintain adipocyte health by promoting M2-like macrophages that act to preserve metabolic function and prevent adiposity and systemic insulin resistance^24, 25^. Consistent with this, whilst their levels in both visceral fat tissues (Figure 3a-d) remain relatively constant throughout ageing in male chow-fed mice, they are progressively reduced by a HCD diet. ES-62 treatment provides significant protection against this decline, particularly in retroperitoneal fat where there is no significant difference in the levels of eosinophils between chow and HCD-ES-62 male mice at either d160 or d500. By contrast, the levels of visceral fat eosinophils were even higher in female HCD-PBS mice at d160 than those found in young lean female mice and these were further boosted, albeit not significantly in retroperitoneal fat, by treatment with ES-62. Although these elevated levels of eosinophils declined to below those observed in chow mice between d340-500, ES-62 still maintained increased eosinophil levels relative to those seen in the gonadal fat of the female HFD-PBS group (Figure 3a-d).

**Figure 3.**
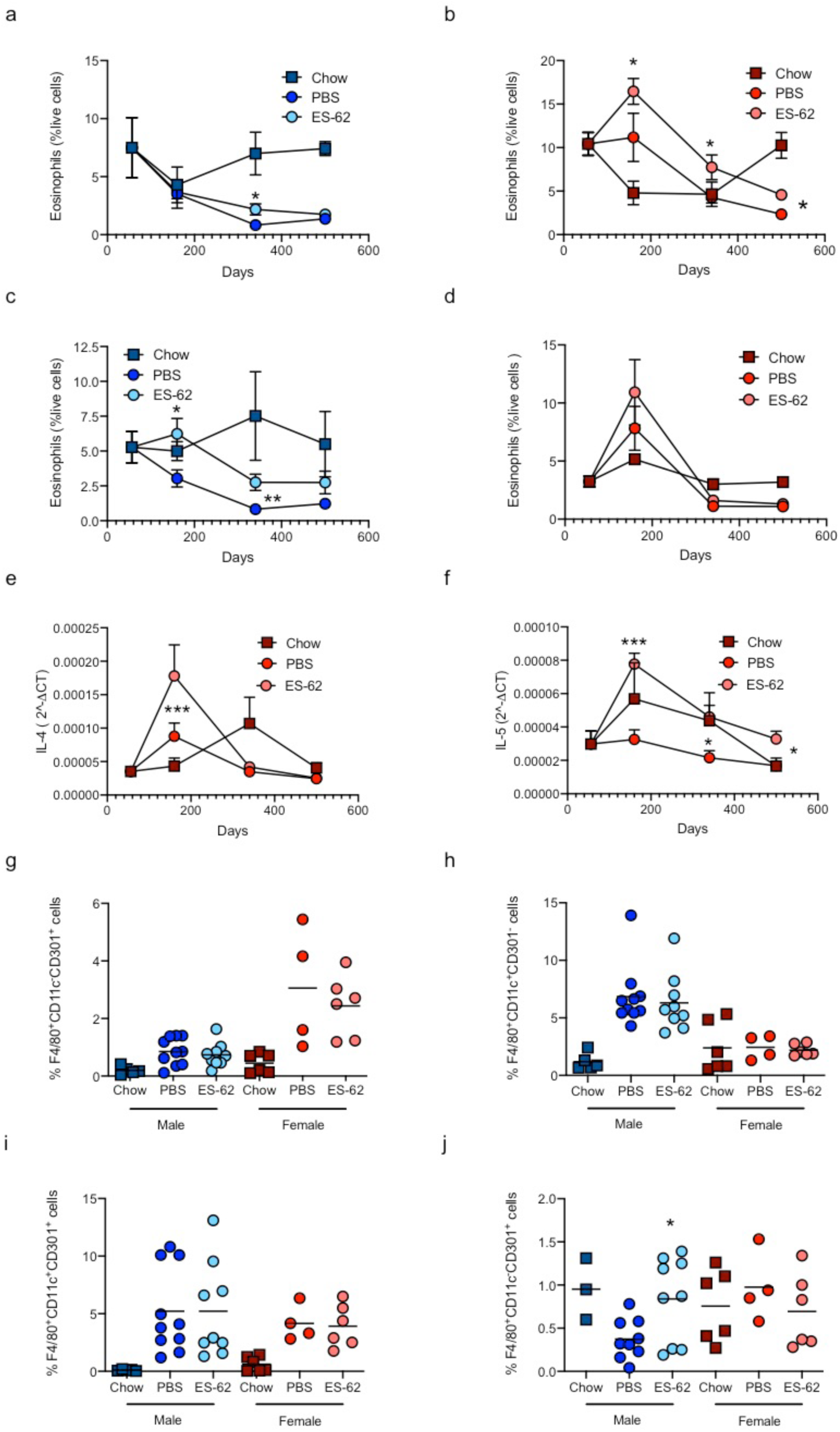
ES-62 promotes eosinophils in visceral fat of HCD-fed mice. The levels of SiglecF^+^ eosinophils (% SVF cells) from gonadal fat of chow- and HCD- (PBS- or ES-62-treated) mice are presented as the mean values ± SEM at each time point where male (a) cohort sizes are: chow - d56, n=5; d160, n=3; d340, n=6; d500, n=5; HCD-PBS - d160, n=10; d340, n=11; d500, n=6; HCD-ES-62 - d160, n=9; d340, n=12; d500, n=6 and female (b) cohort sizes are: chow d56 - n=6; d160, n=6; d340, n=6; d500, n=5; HCD-PBS - d160, n=4; d340, n=11; d500, n=6; HCD-ES-62 - d160, n=6; d340, n=12; d500, n=6. Likewise, levels of eosinophils in retroperitoneal fat are presented as the mean values ± SEM at each time point where male (c) cohort sizes are: chow - d56, n=5; d160, n=3; d340, n=5; d500, n=5; HCD-PBS - d160, n=9; d340, n=10; d500, n=6; HCD-ES-62 - d160, n=9; d340, n=12; d500, n=6 and female (d) cohort sizes are: chow - d56, n=6; d160, n=6; d340, n=6; d500, n=5; HCD-PBS - d160, n=4; d340, n=11; d500, n=6; HCD-ES-62 - d160, n=6; d340, n=12; d500, n=6. qRT-PCR analysis of IL-4 (e) and IL-5 (f) mRNA expression in gonadal fat from female chow- and HCD- (PBS- or ES-62-treated) mice where data are expressed as mean 2^ΔCT values ± SEM of individual mice and the values for each mouse are means of n=3 replicate analyses. Cohort sizes: chow - d56, n=6; d160, n=5; d340, n=5; d500, n=5; HCD-PBS - d160, n=8 (IL-4), n=9 (IL-5) d340, n=11; d500, n=6; HCD-ES-62 - d160, n=9 (IL-4), n=10 (IL-5); d340, n=11; d500, n=6. The levels of F4/80^+^CD11c^-^CD301^+^ (g, j), F4/80^+^CD11c^+^Cd301^-^ (h) and F4/80^+^CD11c^+^Cd301^+^ (i) macrophages (% live SVF cells) in gonadal (g-i) and retroperitoneal (j) fat in male and female chow- and HCD- (PBS- or ES-62-treated) mice in the d160 cohorts are shown. For clarity, only significant differences between the HCD-PBS and HCD-ES-62 cohorts are shown on the figures, where significance is denoted by *p < 0.05, **p < 0.01 and ***p < 0.001.

These sex- and depot-specific effects presumably reflected ES-62 recruitment of differential homeostatic mechanisms capable of counteracting HCD-induced adipocyte dysfunction: in female mice, the increased levels of eosinophils correlated with a rise in IL-4 and IL-5 mRNA expression in the gonadal fat at d160 (Figure 3e & f). HCD-induced changes in adipose metabolic function have been associated with disruption of the phenotypic balance of M1/M2-like adipose tissue macrophages, typically resulting from an increase/recruitment in M1-like (CD11c^+^CD301b^-^) cells, which can be counteracted by the protective actions of eosinophils in promoting an M2 (CD11c^-^CD301b^+^)-like phenotype in an IL-4/5-dependent manner. Our analysis did not reveal a clear ES-62 repolarisation of these macrophage populations in either fat depot from male or female HCD mice during ageing. However, corresponding with the increased eosinophils and type-2 cytokines (IL-4 and IL-5) observed in gonadal tissue from female, but not male, HCD-fed mice at d160, there is a greater increase in M2 (CD11c^-^CD301b^+^)-like macrophages in both PBS- and ES-62-treated female HCD mice than in the male HCD cohorts, which instead displayed a strong rise in M1 (CD11c^+^CD301b^-^)-like macrophages (Figure 3g & h). Nevertheless, it is increasingly evident that M1- and M2-like phenotypes are heterogeneous and that hybrid M1-M2 phenotypes develop during obesity^26, 27^, such as the CD11c^+^CD301b^+^ subset identified in gonadal and subcutaneous fat, depletion of which prevents weight gain (and even promotes weight loss under conditions of obesity) and insulin resistance^28^. Consistent with this, we found CD11c^+^CD301b^+^ macrophages to be increased in gonadal fat (Figure 3i) and, albeit not significantly, in retroperitoneal fat from all male and female HCD-fed groups at d160. Intriguingly, the CD11c^-^CD301b^+^ M2-like population is actually depleted in the retroperitoneal fat of male, but not female, HCD-PBS mice relative to normal chow controls and exposure to ES-62 restores these cells back to the levels found in the male chow mice (Figure 3j). These distinct sex- and fat tissue-specific responses to HCD may potentially contribute to the differential effects of ES-62 on male and female longevity under conditions of diet-induced obesity, especially when considered in the context that increased intra-abdominal visceral fat appears to play an particularly important role in driving obesity-associated pathologies in men, relative to women^29^.

### ES-62 and Adipocyte Dysfunction and Insulin Resistance

Adipocyte hypertrophy has been reported to lead to insulin resistance and liver fibrosis via dysregulation of adipokines like leptin and adiponectin. Whilst leptin normally acts to suppress appetite and adiponectin promotes insulin sensitivity, elevated leptin levels in human obesity appear to reduce adiponectin sensitivity^30, 31^. Consistent with this, whilst serum levels of leptin remained constant in the ageing male and female lean cohorts, despite fat mass increasing, leptin levels rose progressively in ageing male HCD-PBS mice and, quite dramatically between d340 and 500 in female HCD-PBS mice, findings suggestive of leptin resistance: these rises were attenuated in ES-62-treated mice, where leptin levels were not significantly different from those observed in age-matched chow-fed animals (Figure 4a & b). By contrast, the rise in serum adiponectin observed with age in lean male and female mice was not seen in PBS- or ES-62-treated HCD mice (Supplementary Figure 3e & f).

**Figure 4.**
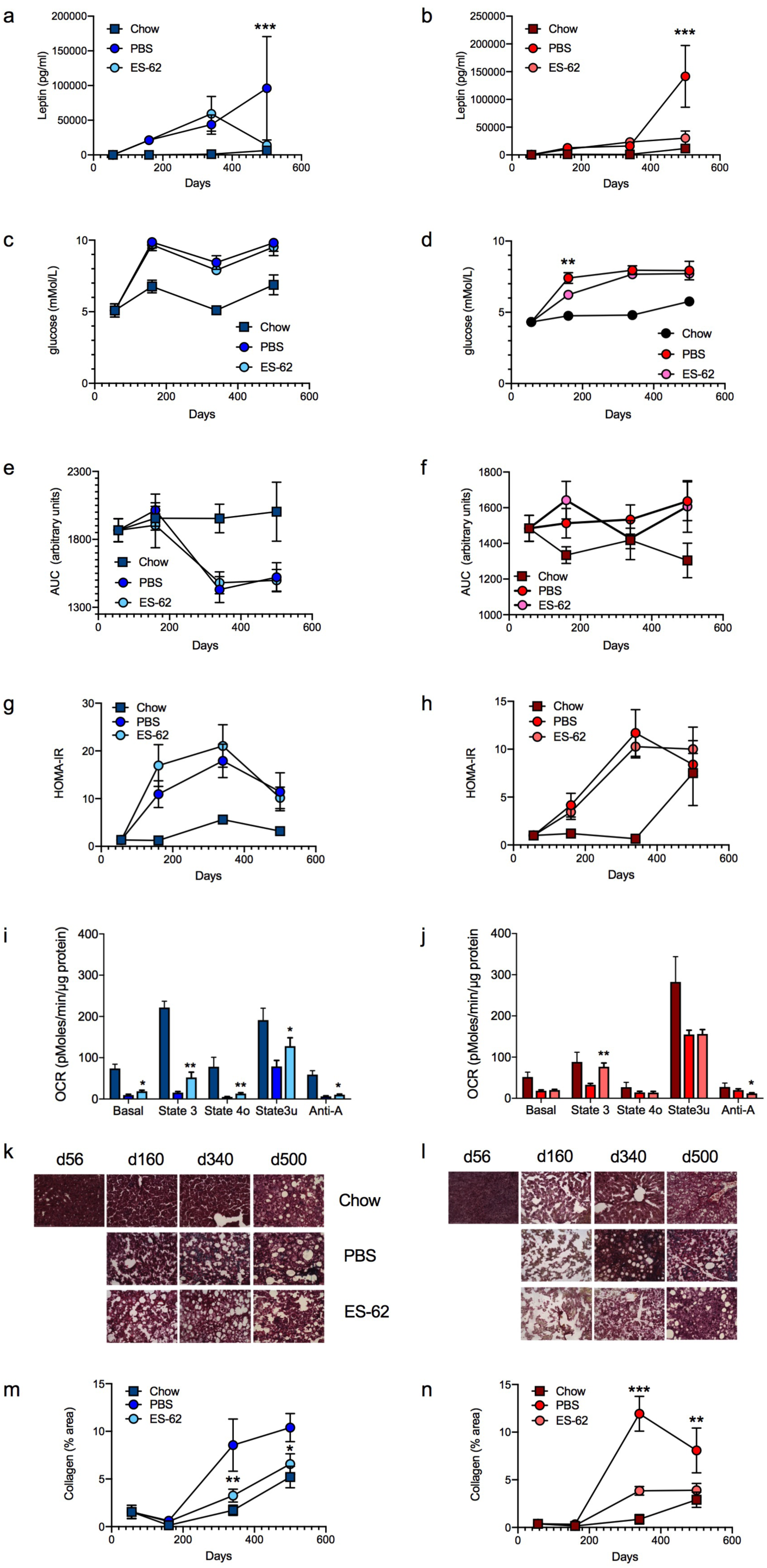
Effect of ES-62 on metabolic function in HCD-fed mice. Mean serum leptin values ± SEM from individual chow- or HFD- (PBS- or ES-62-treated) male (a) and female (b) mice. Measurement of fasting glucose (c, d) and glucose tolerance (GTT-AUC, e, f) were undertaken one week before cull days and presented as mean values ± SEM of individual male (c, e) and female (d, f) mice. Mean HOMA-IR values ± SEM were determined from serum fasting glucose and insulin levels at cull for male (g) and female (h) mice. Cohort sizes: males: chow - d56, n=5 (leptin), 6 (glucose, GTT-AUC, HOMA-IR); d160 and d340, n=6; d500, n=5 (leptin; glucose; GTT-AUC), 4 (HOMA-IR); HCD-PBS - d160, n=9 (leptin; HOMA-IR), 10 (glucose; GTT-AUC); d340, n=8 (leptin), 11 (glucose, GTT-AUC), 9 (HOMA-IR); d500, n=3 (leptin), 6 (glucose, GTT-AUC, HOMA-IR); HCD-ES-62 - d160, n=9 (leptin; HOMA-IR), 10 (glucose; GTT-AUC); d340, n=7 (leptin), 12 (glucose, GTT-AUC), 9 (HOMA-IR); d500, n=3 (leptin), 6 (glucose, GTT-AUC, HOMA-IR) and females: chow - d56, n=6; d160 and d340, n=5; d500, n=3; HFD-PBS - d160, n=9; d340, n=9 (leptin, HOMA-IR), 11 (glucose, GTT-AUC); d500, n=4 (leptin), 6 (glucose, GTT-AUC, HOMA-IR); HCD-ES-62 - d160, n=9 (leptin, HOMA-IR), 10 (glucose, GTT-AUC); d340, n=9 (leptin, HOMA-IR), 12 (glucose, GTT-AUC); d500, n=4 (leptin), 6 (glucose, GTT-AUC, HOMA-IR). Mitochondrial respiration (oxygen consumption rate, OCR) was measured in livers from male (i) and female (j) chow- or HFD-mice at d160. OCR was measured under Basal (substrate alone), State 3 (ADP), State 4 (oligomycin), State 3u (FCCP) and non-mitochondrial (antimycin A plus rotenone; Anti-A) conditions. Data are presented as values ± SEM of individual male (chow n=6; HCD-PBS n=10; HCD-ES-62 n=9) and female (chow n=5; HCD-PBS n=9; HCD-ES-62 n=10) mice. Representative images (scale bar 100 µm) of liver from male (k) and female (l) mice stained with Gömöri’s Trichrome: quantitative analysis of collagen deposition is presented as mean (of triplicate analyses) values ± SEM, where n=5-6 individual male (m) and female (n) mice. Significant differences between the HCD-PBS and HCD-ES-62 cohorts are shown where *p < 0.05, **p < 0.01 and ***p < 0.001.

Consistent with these adipokine effects contributing to insulin resistance, and mirroring human physiology in obesity, serum insulin levels were substantially elevated in female and particularly, male HCD mice: however, this increase was not modulated by ES-62. Of note, ageing (d500) female, but not male, chow-fed mice show elevated serum insulin levels not significantly different to those of their age-matched HCD-fed cohorts (Supplementary Figure 3g & h). Reflecting these changes, increased fasting glucose levels and impaired glucose clearance (as indicated by their glucose tolerance test [GTT] responses) were evident in HCD- but not chow-fed mice by d160 onwards (Figure 4c-f; Supplementary Figure 3i & j). Although ES-62 slightly attenuated the age-associated rise in fasting glucose levels observed in ageing female HCD-PBS mice, it did not protect against the impaired glucose clearance observed following administration of a bolus of glucose to these mice, relative to their chow-fed controls (Figure 4f; Supplementary Figure 3j). This defect in glucose clearance is slight compared to that widely documented for female mice acutely fed a HCD (up to ∼10 weeks), likely reflecting the remodelling of glucose handling reported to result from the increased adipose tissue providing novel glucose sinks in mice fed a chronic HCD^32^. What is striking here however is the dramatic extent of the remodelling of this response in male, relative to female, HCD-fed mice (Figure 4e & f) indicating distinct sex-dependent metabolic responses to obesity. As ES-62 did not significantly modulate glucose tolerance in (either male or female) HCD-fed mice (Figure 4e & f; Supplementary Figure 3i & j), its healthspan-promoting effects appear to be quite distinct from the improved glucose tolerance previously observed in acute mouse models of obesity following infection with helminths, or treatment with their products^33–37^.

Homeostatic model assessment of insulin resistance (HOMA-IR), a test designed to assess IR and pancreatic β-cell function, confirmed that HCD-fed mice, particularly the male cohorts, progressively developed IR and that this was not prevented by ES-62 (Figure 4g & h). Again, we found substantial differences between the male and female cohorts, with the female, but not male, d500 chow-fed group exhibiting HOMA-IR responses similar to their age-matched HCD-fed counterparts (Figure 4g & h). Analysis of pancreatic β-cell function revealed that male HCD-fed mice showed an increase in the size of the pancreatic islets and associated elevated β-cell insulin and α-cell glucagon expression followed by a sharp decline in each of these parameters between d340-500 (Supplementary Figure 4a-h). These data are consistent with the compensatory hyper-production of insulin and glucagon occurring in response to obesity-induced IR, followed by the islet death and pancreatic failure associated with established Type-2 diabetes^30^. The male chow-fed group also showed increases in islet size and insulin and glucagon production at d500 indicative of an emerging ageing-associated IR that is a common feature of ageing in mice and humans^23^. These IR effects were much less pronounced in female mice with the HCD cohort more mirroring the responses of the male chow group. ES-62 had only marginal effects on these pancreatic responses in male HCD-fed mice but appeared, particularly at d160, to convert the female HFD-mice to a phenotype more resembling that of the matched male cohort (Supplementary Figure 4a-h).

### ES-62 promotes liver health

HCD-mediated disruption of adipocyte health and induction of IR consequently impacts on liver function^38^: this is associated with mitochondrial dysfunction and REDOX imbalance, which coupled with inflammation and fat and collagen deposition, can lead to liver steatosis, fibrosis and cancer. Consistent with this, HCD and ageing impaired respiration rates within isolated liver mitochondria of both male and female mice (Figure 4i & j; Supplementary Figure 5a-d). ES-62 partially rescued the HCD-induced impairment in mitochondrial function in male mice at d160 (Figure 4i) but its effects in female mice were much less pronounced, although ES-62 similarly increased ADP-driven respiration rates at 160d (Figure 4j). Such beneficial effects were not observed in the older HCD-fed cohorts (Supplementary Figure 5a-d) and did not reflect protection against the HCD-accelerated decrease in cytochrome C expression (Supplementary Figure 5e & f) previously associated with age-dependent decreases in mitochondrial respiratory capacity^39^.

HCD-exacerbation of mitochondrial dysfunction was also accompanied, particularly in the livers of female mice, by a transient increase in HMOX1 expression (d160) and superoxide dismutase (SOD) and protein carbonylation activities (both d340) in the HCD relative to the chow cohorts, the latter displaying low levels of these REDOX parameters (particularly SOD) across all time-points (Supplementary Figure 5g-l). The ES-62 delay in ageing-associated mitochondrial dysfunction in male HCD-fed mice was not associated with any modulation of these REDOX factors. However, ES-62 further increased HMOX1 expression and of note, decreased the SOD and protein carbonylation activities in female HCD-fed mice, reducing the latter two activities to the levels seen in both PBS- and ES-62-treated male HCD-fed cohorts at d340.

Reflecting the impact of HCD on adipose tissue, there was a dramatic but transient increase in the levels of fatty liver observed by d160 in both male and female HCD-fed mice that then appeared to resolve/remodel, resulting in similar slightly elevated levels of fatty liver being exhibited by the chow and HCD-fed mice in the d500 cohorts (Supplementary Figure 6a-d). Paralleling the dramatic increase in liver fat deposition at d160, transient spikes in inflammatory cytokine (IL-1β and IL-18) and associated NLRP3 expression occurred within livers from HCD-fed mice, particularly of female animals, before returning towards, or below, the stable basal levels observed in their chow counterparts at d500 (Supplementary Figure 6e-j). Reflecting its marginal protective effects on liver steatosis at d500, ES-62 only slightly reduced some of these inflammasome responses at d340 and d500 (Supplementary Figure 6e-j).

Consequent to the spikes in liver steatosis and inflammation, we observed substantial acceleration and enhancement of age-related liver fibrosis (as evidenced by collagen deposition) in both male and female HCD-PBS mice. Critically, exposure to ES-62 ameliorated HCD-induced liver fibrosis in both sexes, maintaining the slow kinetics and low extent of age-related liver fibrosis observed in the chow-fed controls (Figure 4k-n).

### ES-62 acts to maintain gut integrity and prevent inflammation

Emerging evidence from Drosophila suggests that lifespan is limited by gut pathology such that intestinal epithelial barrier integrity appears to be a more effective predictor of mortality than chronological age^40, 41^. Indeed, it has been proposed that the extension of lifespan in response to dietary restriction (DR) in female flies may reflect DR’s ability to reduce their profound gut pathology observed during ageing^42^. We therefore next analysed the effect of ES-62 treatment on HCD-induced gut inflammation, loss of barrier integrity and associated dysbiosis of the microbiome, factors linked not only to the development of visceral fat inflammation, IR and obesity^43, 44^ but also to the acceleration of ageing and shortening of lifespan^45, 46^.

HCD was found to accelerate and exacerbate ileum pathology occurring during ageing of chow-fed mice not only in terms of an overall pathology score (scoring based on degree of cellular infiltration, epithelial erosion and villus atrophy^47^) but also by quantitative analysis of villi length, basal lamina thickness and collagen deposition in both male and female mice (Figure 5a-f; Supplementary Figure 7a-d). ES-62 afforded some protection by slowing the age-associated shortening in villus length in male and, to a lesser (non-significant) extent, female mice (Figure 5e & f). By contrast, analysis of colon tissue showed ES-62 acted to slow the induction of pathology and prevent the changes in crypt morphology (ratio of crypt depth to intercrypt width^47, 48^) in male, but not female, HCD-mice (Figure 5g-l). No effects of ES-62 on colon basal lamina thickness and collagen deposition were observed (Supplementary Figure 7e-h).

**Figure 5.**
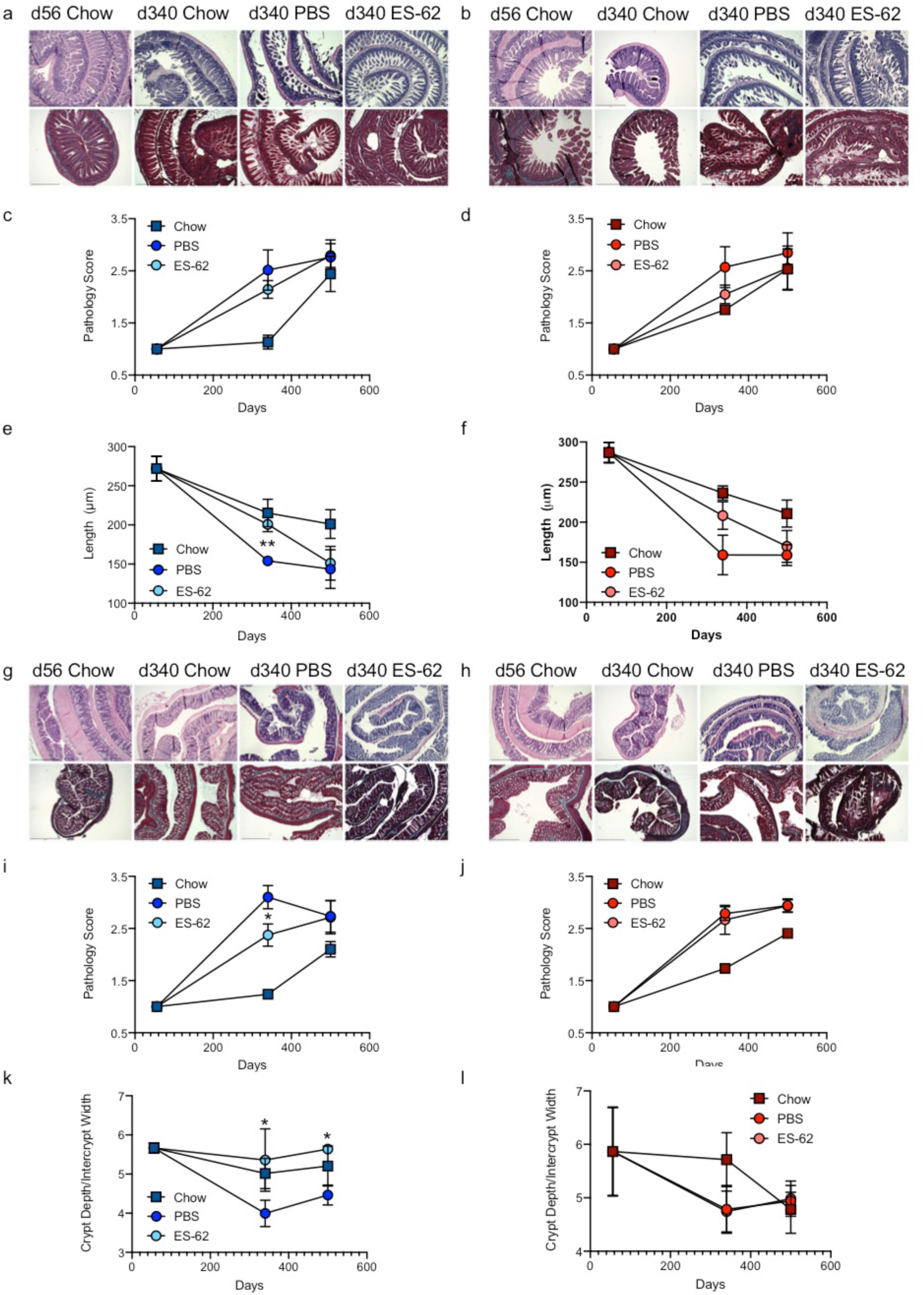
ES-62 protects against gut pathology in HCD-fed mice. Representative images (scale bar 500 µm) of ileum tissue from male (a) and female (b) chow- and HCD- (PBS- or ES-62-treated) d340 mice stained with H & E (upper panels) and Gömöri’s Trichrome and resultant pathology scoring (c, d) and quantitative analysis of ileum villus length (e, f) are shown. Data are presented as the mean values ± SEM where n=4-6 individual male (a, c, e) and female (b, d, f) mice at each time point and the values for each mouse are means derived from n=3 replicate analyses. Representative images (scale bar 500 µm) of colon tissue from male (g) and female (h) chow- and HCD- (PBS- or ES-62-treated) d340 mice stained with H & E (upper panels) and Gömöri’s Trichrome and resultant pathology scoring (i, j) and quantitative analysis of the ratio of crypt depth:intercrypt width (k, l) are shown. Data are presented as the mean values ± SEM where n=4-6 individual male (g, i, k) and female (h, j, l) mice at each time point and the values for each mouse are means derived from n=3 replicate analyses. For clarity, only significant differences between the HCD-PBS and HCD-ES-62 cohorts are shown on the figures, where significance is denoted by *p < 0.05 and **p < 0.01.

Furthermore, reflecting the cellular infiltration and inflammation identified in the pathology scoring, whilst the male mice in the HCD-PBS cohort exhibited increased levels of IL-17 expression in colon relative to their chow-fed counterparts, this was abrogated by exposure to ES-62. By contrast, colonic IL-17 expression progressively increased with ageing in female mice, irrespective of diet or exposure to ES-62 (Figure 6a-c). Age-associated B cells (CD19^+^CD21^-^CD23^-^CD11c^+^ B cells) have been proposed to promote TH1/TH17-mediated inflammation (and autoimmunity) during ageing, although they have also been reported to play roles in protective immunity, particularly with respect to viral infection^49, 50^. Levels of these circulating cells are increased in the spleens of d500 mice (Figure 6d & e), in particular as reported previously, in the female cohorts^49, 50^. Their accumulation is accelerated by HCD and this is not prevented by ES-62: rather, perhaps reflecting their reported relatively high phosphorylcholine (PC)-reactivity, they are increased in both the male and female ES-62-treated HCD-treated cohorts as a likely consequence of chronic exposure to the PC-containing ES-62^4, 5^ (Figure 6f-i). Enhancement of the anti-PC antibody repertoire could possibly serendipitously boost health- and lifespan by helping to protect against bacterial infection in old age^51^ given the increased virulence, morbidity and mortality of (PC-containing) *Streptococcus pneumoniae* and Haemophilus influenza in the elderly^52, 53^. Interestingly, therefore, we have found HCD to reduce the levels of anti-PC antibodies in male and female mice, with this reaching statistical significance with respect to the levels of anti-PC IgG antibodies in male mice relative to the chow-fed counterparts at d340 (*p<0.05).

**Figure 6.**
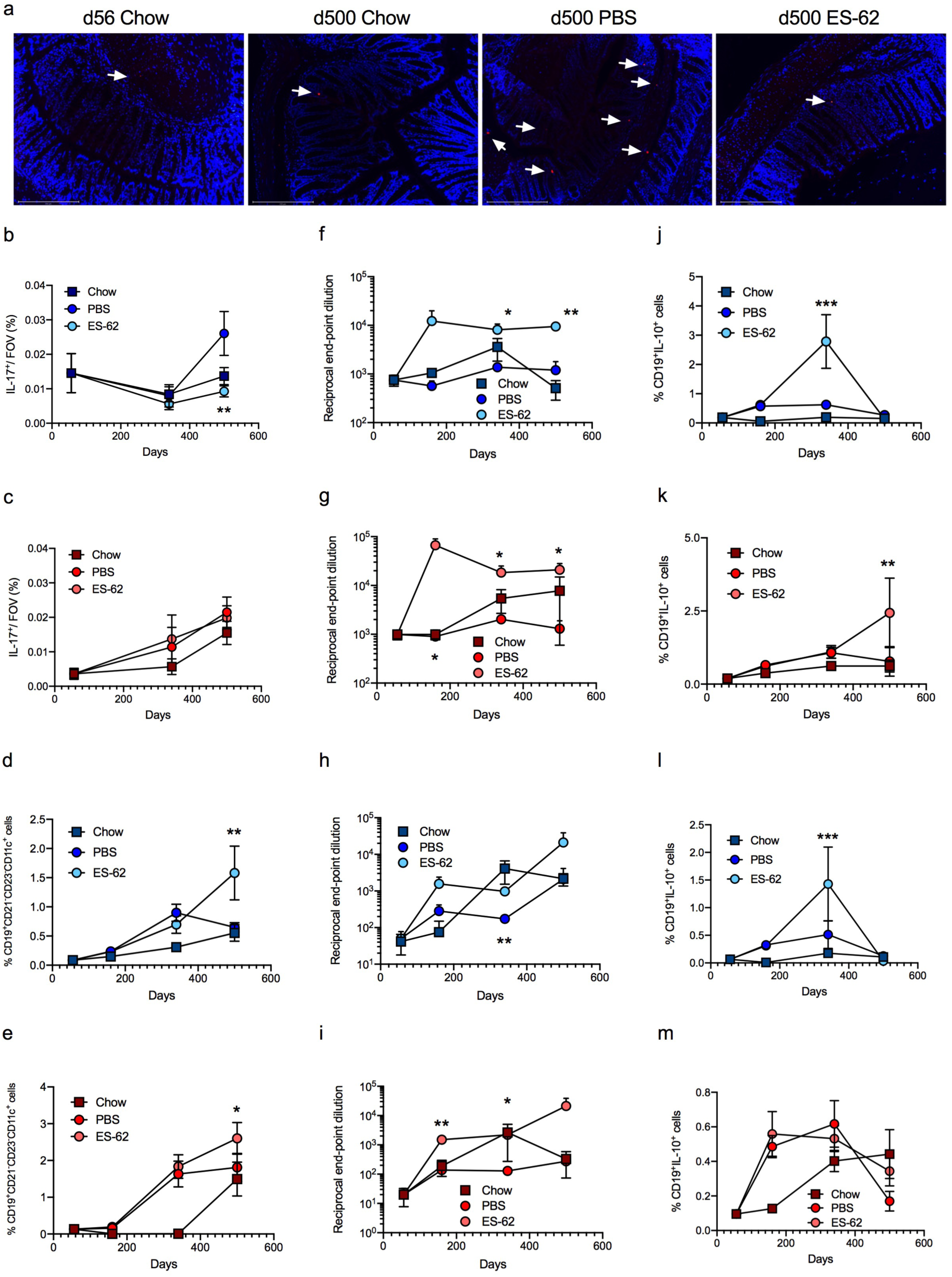
ES-62 modulates B cell responses in HCD mice. Representative images (a; scale bar 200 µm; d500) and quantitative analysis (b, c) of colon IL-17-expression (red, indicated by arrows; DAPI counterstain, blue): data represent mean (of triplicate analyses) values ± SEM, where n=4-6 individual mice per time point. Splenic CD19^+^CD21^-^CD23^-^CD11c^+^ B cells (% live cells) presented as mean values ± SEM where male (d) cohort sizes: chow - d56, n=6; d160, n=6; d340, n=6; d500, n=5; HCD-PBS - d160, n=6; d340, n=7; d500, n=6; HCD-ES-62 - d160, n=6; d340, n=8; d500, n=6 and female (e) cohort sizes: chow - d56, n=5; d160, n=6; d340, n=6; d500, n=3; HCD-PBS - d160, n=9; d340, n=11; d500, n=6; HCD-ES-62 - d160, n=10; d340, n=12; d500, n=6. Reciprocal end-point dilutions of anti-PC IgM (f, g) and IgG (h, i) antibodies as mean values ± SEM where male (f, h) cohort sizes: chow - d56, n=6; d160, n=6; d340, n=6; d500, n=5; HCD-PBS - d160, n=6; d340, n=10 (IgM), 11(IgG); d500, n=6; HCD-ES-62 - d160, n=9; d340, n=11; d500, n=6 and female (g, i) cohort sizes: chow - d56, n=5; d160, n=5 (IgM), 6 (IgG); d340, n=5 (IgM), 6 (IgG); d500, n=5 (IgM), 3 (IgG); HCD-PBS - d160, n=7(IgM), 8 (IgG); d340, n=8 (IgM), 7 (IgG); d500, n=6; HCD-ES-62 - d160, n=8; d340, n=10 (IgM), 11 (IgG); d500, n=6. Splenic (j, k) and MLN (l, m) Bregs (CD19^+^IL-10^+^; % live cells) as mean values ± SEM where male (j, l) cohort sizes: chow - d56, n=6; d160, n=6; d340, n=6; d500, n=5; HCD-PBS - d160, n=10; d340, n=11 (splenic), 7 (MLN); d500, n=6; HCD-ES-62 - d160, n=10; d340, n=12 (splenic), 8 (MLN); d500, n=6 and female (k, m) cohort sizes: chow - d56, n=5 (splenic) 6 (MLN); d160, n=6; d340, n=5 (splenic), (6 (MLN); d500, n=5 (splenic), 4 (MLN); HCD-PBS - d160, n=9; d340, n=10; d500, n=6 (splenic), 5 (MLN); HCD-ES-62 - d160, n=10 (splenic), 9 (MLN); d340, n=12 (splenic), 10 (MLN); d500, n=6. Significant differences between HCD-PBS and HCD-ES-62 cohorts are shown where *p < 0.05, **p < 0.01 and ***p < 0.001. (350 max 350).

Nevertheless, analysis of CD19^+^IL-10^+^Bregs (Figure 6j-m), cells that can act to resolve pathogenic inflammation^54^ including that associated with perturbation of the gut microbiota^55^ and obesity^56^, showed that exposure to ES-62 increased their levels in the spleens of male (d340) and female (d500) HCD-fed mice (Figure 6j & k). Interestingly, and presumably via a homeostatic mechanism to control the gut pathology emerging in d500 chow-fed mice, we find increased levels of CD19^+^IL- 10^+^Bregs in the MLN of female chow-fed mice by d340-500. Presumably reflecting the HCD-accelerated gut pathology, Breg levels are elevated in HCD-fed female mice at d160-340 before those in the PBS, but not ES-62, groups decline sharply by d500. By contrast, whilst there is little change in the levels of CD19^+^IL-10^+^ B cells in MLNs of chow- or PBS-treated HCD-fed male mice, exposure to ES-62 substantially increases the levels of these cells at d340 (Figure 6l & m).

### ES-62 maintains gut microbiota homeostasis during HCD-induced ageing

Changes in the microbiome contribute to the loss of intestinal barrier integrity^43, 44, 57^ during ageing and impact directly on obesity and longevity^45, 46^. We have recently shown that the protection afforded by ES-62 against inflammatory arthritis is associated with normalisation of the gut microbiome and stabilisation of gut barrier integrity that prevents generation of pathogenic TH17-mediated inflammation and allows homeostatic resolution of inflammation by IL-10^+^ Bregs in the collagen-induced arthritis model of rheumatoid arthritis in male mice^12^. Thus, we investigated whether ES-62 acted to normalise the ageing microbiota and prevent the dysbiosis driving loss of barrier integrity, by metagenomic analysis of faecal (ileum plus colon) material, with samples from individual mice pooled to generate a representative cohort phenotype. Whilst generally stable in healthy adults, the gut microbiome can undergo dynamic changes particularly in early and late life and during inflammatory and metabolic disorders^58^. Consistent with this, we found profound differences at the phylum level in the microbiota not only amongst young and ageing male and female chow-fed mice, but also to a lesser extent between aged-matched chow- and HCD-PBS mice, with ES-62 acting to normalise the impact of HCD back towards the chow profile, particularly in male mice (Figure 7a, upper panels; day 340 data shown). There was a striking loss of Verrucomicrobia, particularly in female mice (Figure 7a, lower panels), which exhibited unexpectedly high levels of these bacteria in the young (d56) control cohort and the d340 female chow-fed group. Verrucomicrobia (primarily of the genus Akkermansia) are mucus-degrading commensals that have been reported to be a marker of gut health due to their ability to promote gut barrier integrity, Treg generation and insulin sensitivity^59, 60^: thus, these findings may suggest, as with flies^42^, that female C57BL/6J mice may be particularly vulnerable to (HCD-induced) gut pathology during ageing. The failure of ES-62 to restore these commensals, and promote Treg responses in general^4^, may go some way to explaining why the helminth product does not extend lifespan in female HCD-fed mice but only in the male cohorts, which given their lower levels of Verrucomicrobia may be innately less susceptible to the pathological consequences of gut dysfunction. Certainly, there is sexual dimorphism in (microbiota-driven) immune responses and consequently, disease susceptibility with notably the stronger immune responses to infection evident in females predisposing them to chronic inflammatory disorders^61, 62^.

**Figure 7.**
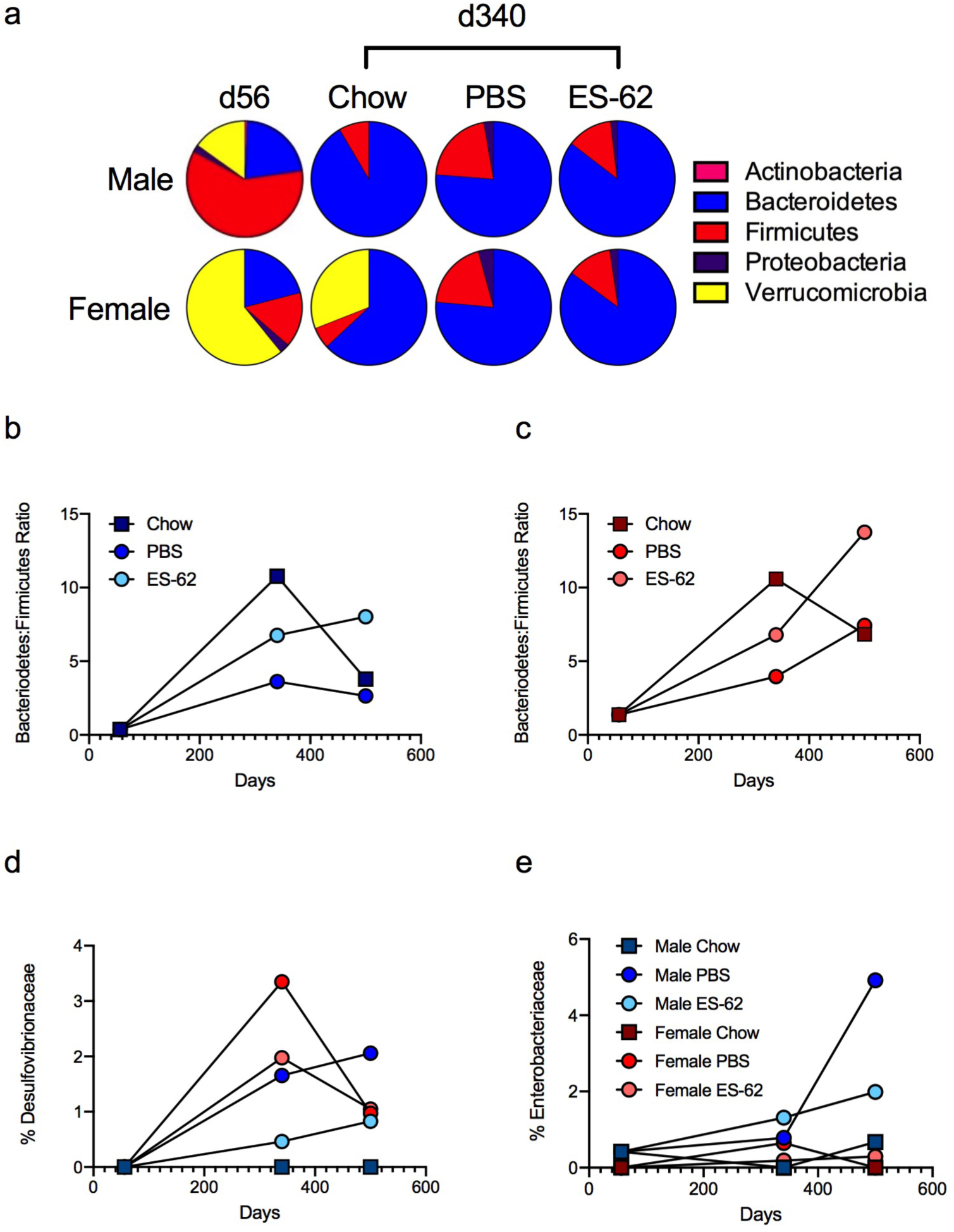
ES-62 acts to normalise the gut microbiome in HCD-fed mice. The composition of bacterial phyla present in gut (ileum plus colon) faecal matter (a) of male and female chow- (d56 and d340) and HCD- (PBS- or ES-62-treated; d340) mice presenting proportion values as pie charts using pooled samples to represent each cohort. The Bacteroidetes:Firmicutes ratio (b, c) and levels of Desulfovibrionaceae (d) and Enterobacteriaceae (e) present in gut (ileum plus colon) faecal matter of male (a, b, d, e) and female (a, c, d, e) chow- and HCD- (PBS or ES-62-treated) mice (each determined as proportion values) using pooled samples to represent each cohort at the indicated time-points are shown.

Typically, whilst the ratio of Bacteriodetes:Firmicutes increases from birth through to adulthood, it subsequently remains relatively stable in healthy individuals before decreasing in old age^63^. However, obese adult humans show a decreased Bacteroidetes:Firmicutes ratio relative to aged-matched lean individuals^46^. Reflecting this, whilst the proportions of Bacteriodetes:Firmicutes in the faecal (ileum plus colon) matter of young (d56) lean male and female mice are low, mid-life adult (day 340) chow-fed male and female mice exhibit an (equivalent) much higher Bacteriodetes:Firmicutes ratio which is reduced by some 65% in both sexes of HCD-PBS mice (Figure 7b & c). As expected, the Bacteroidetes:Firmicutes ratio in both sexes of the chow-fed mice declined during ageing but this was more pronounced in the male relative to the female d500 cohorts (by ∼65% and ∼35%, respectively), whilst perhaps surprisingly, the female HCD-PBS group slowly increased its Bacteroidetes:Firmicutes ratio across the cross-sectional timepoints. Exposure to ES-62 protects against the HCD-induced decreases in the Bacteroidetes:Firmicutes ratio in both male and female mice and indeed, maintains, or even increases, their proportional abundance in the d500 HCD-fed cohorts (male, ∼75%; female, ∼130%, relative to the mid-life adult d340 chow-fed animals; Figure 7b & c).

Increases in the proportions of Proteobacteria relative to Firmicutes have been associated with loss of gut barrier integrity and proposed to provide signatures of gut inflammatory disorders (e.g. inflammatory bowel disease) and ageing^58^. Interestingly, the Desulfovibrionaceae family of δ-Proteobacteria negatively correlate with longevity and have been shown to be reduced following DR in male C57BL/6J mice^64^, whilst the Enterobacteriaceae (γ-Proteobacteria) are positively correlated with ageing-induced frailty in flies^57^. Perhaps critically therefore, whilst we found that HCD-feeding induced ageing-associated increases in the Desulfovibrionaceae family of δ-Proteobacteria in male and particularly, female mice, these increases were inhibited by exposure to ES-62 although in the case of female HCD mice, this reduction was only generally to around the level found in male HCD-PBS mice at d340 (Figure 7d). Indeed, this male sex protective bias against pathogenic Proteobacteria was even more strikingly reflected in the finding that HCD induces a strong enrichment of the Enterobacteriaceae in ageing male but not female mice (d500) and that this outgrowth is dramatically reduced by exposure to ES-62 (Figure 7e). Collectively, these data suggest that ES-62 acts to broadly normalise gut microbiome changes associated with ageing and loss of intestinal barrier integrity and, that this is more evident in male HCD-fed mice, suggest that it may be a contributor to protection against HCD-induced mortality otherwise resulting from the consequent dysregulation of immunometabolic pathways.

### Mathematical modelling of ageing “signatures” in chow- and HCD-fed mice

Our survival and cross-sectional intervention studies have highlighted a number of differential age-, sex-, diet- and/or ES-62-treatment responses in ageing mice that underline the need for identification of precise and relevant biomarkers to identify sites of potential intervention and appropriately stratified target cohorts. To begin to address this, we have subjected our multidimensional data sets (113 pathophysiological, immunological and metabolic variables assayed on individual mice in this study) to unsupervised mathematical modelling and statistical analysis to identify the key (differential) signatures associated with ES-62-mediated promotion of healthspan in male and female HCD-fed mice. Firstly, we validated our data set analysis by identifying the features most predictive of HCD-fed mice (Figure 8a) and as expected, this revealed (with >95 % accuracy) increases in adiposity and liver damage as well as adipokine responses associated with IR. Examination of the sex differences in response to the HCD highlighted the inflammatory bias (Figure 8b) of female mice demonstrated in the above experimental studies. Reflecting this bias, prediction of the healthspan features most robustly associated with ES-62-treatment of female HCD-fed mice (Figure 8c) revealed modulation of a panel of almost exclusively, inflammatory markers (anti-PC antibodies, visceral fat eosinophils and type-1 [IL-1β, IL-18] and type-2, [IL-5, IL-10 and IL-13] cytokines). By contrast, whilst ES-62 treatment of male HCD-fed mice (Figure 8d) was also associated with modulation of inflammatory markers, it additionally targeted signatures associated with gut integrity (ileum villus length, colon crypt: intercrypt ratio) and metabolic tissue function (fasting glucose, gonadal adipocyte size, retroperitoneal fat UCP1).

**Figure 8.**
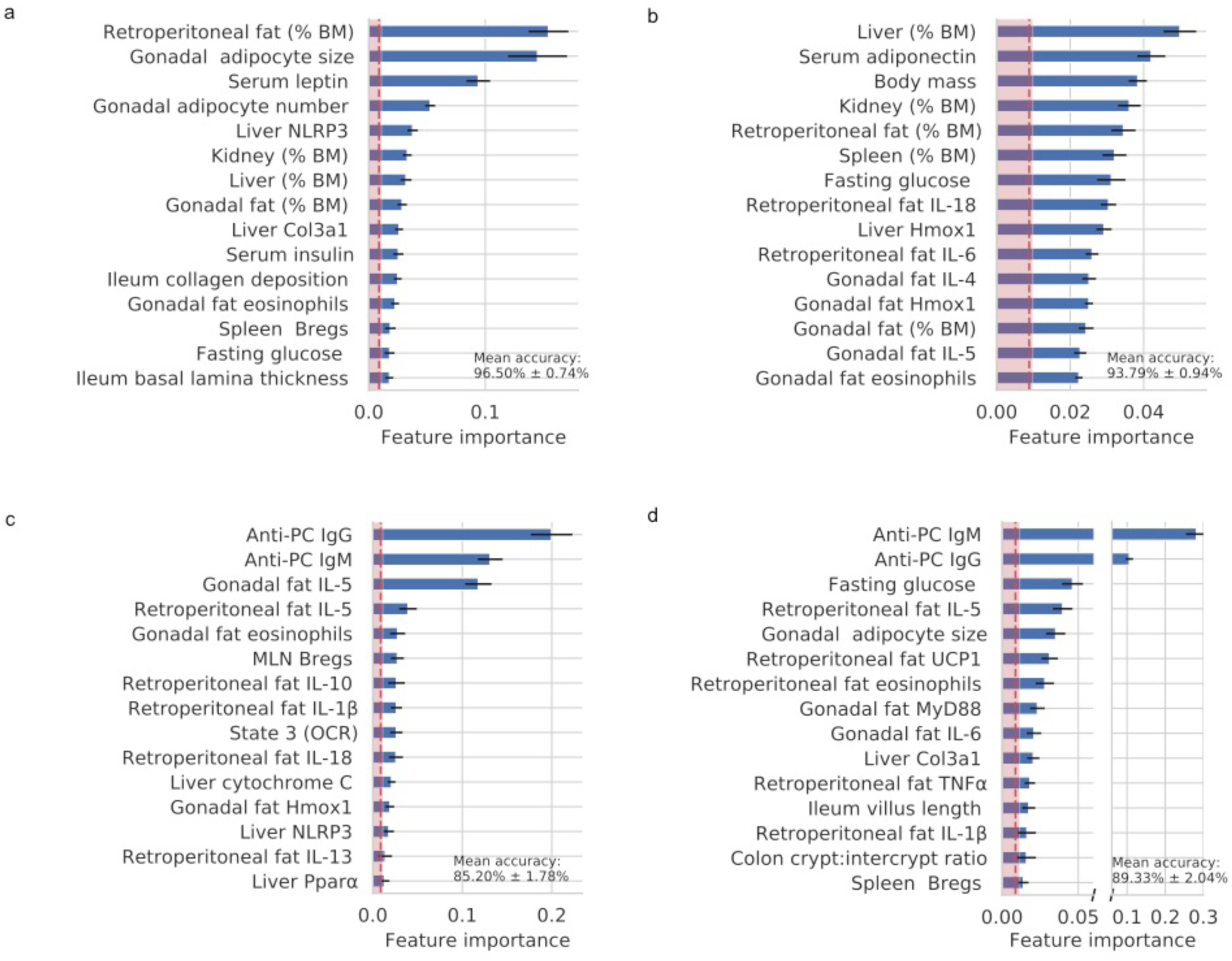
Mathematical modelling of ES-62 healthspan targets. As described in Methods, to identify the pathophysiological, metabolic and immunological variables most robustly associated with ES-62 treatment, sex, and diet, each of the associated mouse cohorts were treated as classes in supervised machine learning classification. The top predictors of (a) HCD-feeding (average (96.50% accuracy of classification), (b) sex-biased effects of HCD-feeding (average 93.79% accuracy of classification) and features most robustly indicative of ES-62-treatment of (c) female (average 85.20% accuracy of classification) and (d) male (average 89.33% accuracy of classification) HCD-fed mice are shown. Each bar represents the mean F score of the corresponding feature, horizontal black lines represent standard errors of the mean, the vertical dotted red line is the global mean F score of all features, and the shaded area is ≤ 1 standard error of the global F score mean. Features considered robustly associated with treatment are those that exceed the shaded area.

## Discussion

Perhaps not surprisingly, the features most robustly predictive of ES-62 treatment in either sex by the machine learning analysis are anti-PC IgM and IgG antibody responses. Of note it has been reported in a number of studies that, in addition to providing protective immunity against *S. pneumonia* and *H. influenza*, anti-PC antibodies, particularly those of the IgM isotype, can protect against development of inflammatory conditions like atherosclerosis^65^. Although the mechanisms involved are not fully understood, it raises the possibility that such antibodies may contribute to the protective effects of ES-62 against HCD-ageing witnessed in the current study. However, although ES-62 protects against lupus-accelerated atherosclerosis, the idiotype of anti-PC antibodies produced is distinct from the T15 idiotype^66^, which appears to be the main idiotype associated with protection against atherosclerosis^65^ and pneumococcal infection^51^. Moreover, PC-based small molecule analogues of ES-62, which mirror the parent molecule in protecting against disease development in models of arthritis, lupus and asthma, are non-immunogenic^4^. Thus, although not ruling out a role for anti-PC antibodies, these findings suggest that other actions of ES-62 likely contribute to the increase in healthspan observed in the current study. Certainly, the observation that IgM anti-PC antibody responses are, if anything, even higher in female mice argues against a role for them in promoting male lifespan. Indeed, taking all of our data and analysis into consideration, prominent amongst factors that contribute to this, may be the improvement in gut and adipose tissue function observed in male ES-62-HCD-mice.

It is worth emphasizing that our study is unique in that we have investigated the effect of a defined parasitic worm product, ES-62, on obesity-accelerated ageing essentially throughout the entire lifespan in a mouse model. It is perhaps remarkable that, when administered only weekly and with low dosage, ES-62 improves multiple aspects of healthspan albeit differentially in male and female HCD-fed mice and substantially increases median lifespan in male mice. Furthermore, an additional outcome of the study has been the revelation of how strongly each of diet and sex can differentially impact on pathophysiological, immunological and metabolic parameters of ageing. Our principal finding however is that ES-62 has provided candidate gut, adipose and liver tissue signatures that merit further exploration as key targets for therapeutic intervention to improve healthspan during ageing and increase lifespan.

## Methods

### Mouse husbandry: longevity and cross-sectional healthspan studies

Male and female C57BL/6J mice (Envigo, UK) were housed in the Central Research Facility (University of Glasgow, Scotland) and maintained at 22°C under a 12-hr light/dark cycle. All procedures were undertaken under a UK Home Office Project Licence (60/4504), following the “principles of laboratory animal care” (NIH Publication No. 86-23, revised 1985) and in accordance with local ethical committee guidelines. Lifespan (survival study) cohort mice (all groups n=24) were housed in same sex groups of 2 to 4 under specific pathogen-free conditions within individually ventilated cages (Techniplast, UK). Mice had *ad libitum* access to water and normal chow (CRM-P; SDS, UK; Oil, 3.36%; Protein 18.35%; Fibre, 4.23%: Sugar 3.9%; Atwater fuel energy from Oil, 9.08%; Protein, 22.03%: Carbohydrate, 68.9%) plus 150 ppm Fenbendazole, until 10 weeks of age when the diet was switched to Western Diet RD, here denoted high calorie diet (HCD; Fat, 21.4%; Protein, 17.5; Fibre, 3.5%; Sucrose 33%; Atwater fuel energy from Fat, 42%; Protein, 15%: Carbohydrate, 43%) plus 150 ppm Fenbendazole. Mice were administered PBS or purified ES-62 (1 μg) weekly via the subcutaneous route from 9 weeks old. Endotoxin-free ES-62 was purified from spent culture medium as described in detail previously^67^. Individuals in the lifespan study were monitored daily, weighed twice-weekly and analysed for grip strength monthly, but otherwise left undisturbed until they died. A pre-weighed amount of food was administered via wire bar lid food hopper. The average consumption/mouse/day was calculated at the end of a 72-hour monitoring period monthly. Grip strength monitoring was carried out as per manufacturer’s guidelines (Ugo Basile®, Italy) using a Gripometer, measuring the peak grip force (GF) of the front limbs. The mouse was lowered towards a T-shaped grasp bar, once the animal gripped the bar it was gently pulled away and the peak amplifier recorded the amount of resistance. An average of three measurements was described as the mean GF.

Survival was assessed from 48 female (HCD-PBS, n=24; HCD-ES-62, n=24) and 48 male (HCD-PBS, n=24; HCD-ES-62, n=24) mice, with all animals dead by the time of this report. Kaplan-Meier survival curves were constructed using known birth and death dates, with differences between groups evaluated using the log-rank test. If death appeared imminent (as assessed by our humane end-points) mice were weighed, euthanized and examined for macroscopic pathological changes using previously described protocols^68, 69^ with this date considered date of death.

Additional cross-sectional healthspan cohorts of mice were sacrificed at 56 (Chow, n=6), 160 (HCD, n=10/group; Chow, n=6), 340 (HCD, n=12/group; Chow, n=6) and 500 (HCD, n=6/group; Chow, n=5) days of age and tissues were harvested for processing. At sacrifice, blood was collected and the serum isolated and stored at −20°C in endotoxin-free Eppendorf tubes. Liver was immediately removed post-mortem and mitochondria prepared for Seahorse analysis^70^. Blood, spleen, mesenteric lymph nodes MLN) and visceral (gonadal: epididymal and parametrial fat pads, and retroperitoneal: dorsal fat pad directly behind the kidneys and attached to the peritoneum proposed as relevant models for human intra-abdominal adipose tissue^71, 72^) fat tissue were prepared for flow cytometry analysis. Ileum and colon faecal contents and liver and visceral adipose tissues were collected in sterile endotoxin-free Eppendorf tubes, snap frozen and stored at −80°C. In addition, all tissue samples were fixed in 10% neutral buffered formalin, and embedded in either paraffin or flash frozen in OCT and stored at −x80°C for histological analysis.

### Tissue processing, histology, and immunofluorescence

Paraffin embedded tissues were sectioned at 5-6 μm and OCT tissues were cryosectioned (Thermoscientific, UK) at 8-10 µm thickness between −14 to −30°C. All tissues were haematoxylin and eosin (H&E) stained, gut and liver tissue sections were also stained with Gömöri’s Trichrome using previously described methods^12^. Prior to immunofluorescence (IF) staining, antigen retrieval (10 mM citrate buffer, pH 6) was performed and the sections were avidin/biotin blocked (Vectorlabs, UK). Briefly, pancreatic insulin and glucagon were stained with rabbit anti-insulin (1/200 dilution: Catalogue number Ab181547 Abcam, UK) and mouse anti-glucagon antibodies (1/200 dilution: Catalogue number Ab10988 Abcam, UK) overnight, followed by goat anti-rabbit IgG Alexafluor 488 (1/400 dilution: Abcam, UK) and goat anti-mouse IgG Alexafluor 647 (1/400 dilution: Catalogue A28181 Invitrogen, UK). IL-17 was detected using goat anti-mouse IL-17 (1/100 dilution, Catalogue number Af-421-na R&D Systems, UK) followed by biotinylated rabbit anti-goat IgG (1/200 dilution; Catalogue number 31732 Invitrogen, UK) with streptavidin Alexafluor 647 (1/200; dilution Catalogue number S21374 Invitrogen, UK). IF sections were mounted with Vectashield with DAPI (Vectorlabs, UK) and images were acquired with an EVOS FL Auto 2 system (Thermofisher, UK).

### Image quantification and analysis

Images were quantified with Volocity (PerkinElmer, USA) or ImageJ Software. The total pancreatic islet number and area (pixels) were quantified and total IF-labelled glucagon and insulin fluorescence (pixels) identified and thresholded across the entire field of view (FOV) using set intensity parameters. Liver fat droplet deposition was calculated as a percentage of thresholded “white area” relative to the total area of liver tissue in FOV. Adipose cell nuclei were identified, thresholded and counted using set intensity parameters and the particle analyser plugin. Adipocount software (CSBIO) was used to quantify the number and area of individual adipocytes in the field of view (FOV). In ileum and colon tissue basal lamina width, villi length (from crypt to tip) or crypt depth was measured in three separate areas in the FOV and averaged. Collagen deposition was quantified by identifying percentage green/blue staining in FOV using the ColourDeconvolution plugin. A gut tissue pathology scoring protocol was developed from the methods described by Erben *et al*.^47^. Images were given an unbiased score for each described parameter. The overall pathology score was calculated (average of cellular infiltration-spatial, epithelial erosion and villus/crypt atrophy).

### Glucose tolerance testing and HOMA-IR

Mice were fasted overnight prior to testing. Throughout the procedure blood samples were obtained via caudal vein venesection and measured using an accu-chek performa (Roche, UK) or OneTouch Ultra (Lifescan, UK) glucometer. A fasting glucose (mmol/L) measurement was taken prior to the intraperitoneal administration of 20% glucose solution (dosage, 2g glucose/kg body mass) and subsequent glucose tolerance measurements were recorded at 15, 30, 60 and 120 minutes post injection, with the glucose tolerance expressed as the area under the curve (AUC) over this 120 minute period^73^. Serum insulin, leptin and adiponectin were assessed using the mouse metabolic and adiponectin MSD kits as per manufacturer’s guidelines (Meso Scale Diagnostics, USA). Insulin resistance was estimated using the homeostasis model assessment for insulin resistance (HOMA-IR) index using the formula described previously^74^, HOMA-IR index = [fasting glucose (mmol/L) x fasting insulin (mIU/L)]/22.5).

### Flow cytometry

Cells from spleen, MLN, whole blood and the stromal vascular fraction (SVF) of gonadal and retroperitoneal adipose tissues were suspended in FACS buffer (2.5% BSA; 0.5 mM EDTA, in PBS) following red blood cell-lysis (eBioscience, UK). Adipose tissue was digested with 1 mg/ml collagenase type II (Sigma, UK) and passed through 100 μM cell filters to generate a single cell suspension. Cells were washed in FACS buffer and incubated with Fc block (Biolegend, UK) before staining with the relevant antibodies/streptavidin conjugates, all of which were purchased from BioLegend, UK unless indicated otherwise. For eosinophils, cells were analysed for SiglecF (PE: Catalogue number 552126 BD Bioscience, UK) expression and for macrophage phenotypes, F4/80 (Biotin: Catalogue number 123105 with SA-ef780), CD301b (APC: Catalogue number 146813) or CD206 (APC: Catalogue number 141707) and CD11c (PeCy7: Catalogue number 117318) expression was determined. Whole blood was red-cell lysed, washed in FACS buffer before incubation with Fc block and the relevant antibodies. Lymphocytes were incubated with antibodies specific for mouse CD3 (FITC or PE: Catalogue number 100203 or 100205), CD4 (PE: Catalogue number 100407 or APC-ef780: Catalogue number 47-0042-82 eBioscience, UK), CD8 (PeCy7: Catalogue number 25-0081-82, eBioscience, UK), CD45RB (FITC: Catalogue number 103305 or APC-ef780: Catalogue number 47-0451-82, Invitrogen, UK) and CD44 (PerCP: Catalogue number 103036). Lymphocytes from the spleen and MLNs were labelled with antibodies specific for mouse CD19 (AF700: Catalogue number 115527), CD23 (PeCy7 or AF488: Catalogue number 101613 or 101609), CD21 (PE: Catalogue number 123409), CD11c (FITC or PeCy7: Catalogue number 117305 or 117318), IgD (PerCP/Cy5.5: Catalogue number 405709), IgM (Biotin: Catalogue number 406503; Strepavidin Bv450), IL-10 (PE: Catalogue number 505007). Fixable viability stain (APC-ef780 or and v450: Catalogue number 65-0865-14 or 65-0863-14; Invitrogen, UK) was used to select for live cells and, for analysis of IL-10^+^ regulatory B cells (Bregs), lymphocytes were stimulated with PMA, ionomycin, Brefeldin A and LPS (Sigma, UK) as described previously^12^. Data were acquired using a BD LSRII flow cytometer and populations were gated using isotype and fluorescence minus one (FMO) controls using FlowJo, LLC analysis software (Tree Star/BD) as described previously^12^. Exemplar gating strategies are presented (Supplementary Figure 8).

### Serum antibody ELISA

Anti-PC-BSA IgG and IgM antibodies in serum^67^ were quantified using a reciprocal end point dilution method. Briefly, high-binding 96 well ELISA plates were coated with PC-BSA overnight at 4°C before washing and blocking with 1% BSA/PBS. Serum was initially diluted 1:100 and then serially diluted three-fold until 1:218700, incubated with HRP-conjugated goat anti-mouse IgG or IgM (1:10,000; 1:6,000 in 10% FBS/PBS) prior to developing with TMB, stopping with 2M sulphuric acid and read at an optical density of 450 nm.

### Seahorse XF assay and REDOX parameters

The XF Mitostress assay (Agilent Technologies, UK) was performed as previously described^70^. Briefly, hepatic mitochondria were isolated and added to an XF culture plate at a concentration of 10 μg per well. A pre-soaked eXF cartridge was prepared^75^, the culture plate loaded into an XF24 Analyser (Agilent Technologies, UK) and the assay initiated. Oxygen consumption rate (OCR) was measured in substrate alone (10 mM pyruvate, 2 mM malate), during state 3 (ADP [4 mM]), state 4 (oligomycin [2.5 mg/ml]), state 3u (FCCP (carbonyl cyanide 4-(trifluoromethoxy) phenylhydrazone) [4 mM]), and lastly antimycin A with rotenone [40 mM]. All reagents were sourced from Sigma, UK. Superoxide dismutase (SOD) and protein carbonyl activities were quantified in liver tissue homogenates using Total SOD activity assay and Carbonyl assay kits, respectively in line with the manufacturer’s guidelines (Cayman Chemical Company, Estonia).

### qRT-PCR

mRNA was extracted from liver and gonadal and retroperitoneal adipose tissue using the RNeasy Lipid Tissue mini kit (Qiagen, Germany), or tissue was lysed in QIAzol prior to mRNA extraction using the DNA away RNA extraction kit (NBS Biologicals, UK). mRNA was transcribed into cDNA using the High Capacity cDNA Reverse Transcriptase kit (Applied Biosystems, Life Technology, UK) for use with Applied Biosystems Quant Studio 7 and KiCqStart→ qPCR ready mix (Sigma-Aldrich) and KiCqStart™ Primers. Data were normalized to the housekeeping gene β-actin to obtain the ΔCT measurements that were used to calculate 2^-ΔCT values. Primer sequences were β-actin (forward - GATGTATGAAGGCTTTGGTC, reverse - TGTGCACTTTTATTGGTCTC), IL-1β (forward - GTGATATTCTCCATGAGCTTTG, reverse - TCTTCTTTGGGTATTGCTTG), IL-4 (forward – CTGGATTCATCGATA AGCTG, reverse - TTTGCATGATGCTCTTTAGG), IL-5 (forward – CCCTACTCATA AAAATCACCAG, reverse - TTGGAATAGCATTTCCACAG), IL-18 (forward – AAAT GGAGACCTGGAATCAG, reverse - CCTCTTACTTCACTGTCTTTG), cytochrome C (forward - CCGGAACGAATTAAAAATGG, reverse –TCTGTGTAAGAGAATC CAGC), HMOX-1 (forward - CATGAAGAACTTTCAGAAGGG, reverse – TAGATATG GTACAAGGAAGCC) and NLRP3 (forward - GATGCTGGAATTAGACAACTG, reverse - GTACATTTCACCCAACTGTAG).

### Metagenomics

Genomic DNA was extracted from the ileum and colon faecal matter using the QIAamp DNA Stool Mini Kit (Qiagen). Colon and ileum DNA were combined and samples were pooled on a group basis for shotgun metagenomic analysis using the Ion Torrent PGM™ platform. Pooled DNA (100 ng) was fragmented and barcoded using the NEBNext→ Fast DNA Fragmentation and Library Prep Set for Ion Torrent™ (NEB Inc, UK) and IonXpress Barcode Adapters kit (ThermoFisher Scientific) respectively. The quality and quantity of all barcoded DNA libraries were analysed using the High Sensitivity DNA analysis kit (Agilent, UK) on the 2100 Bioanalyzer Instrument and Qubit Fluorometer (ThermoFisher Scientific). Samples were prepared using the Ion PGM™ Hi-Q™ View OT2 and Ion PGM™ Hi-Q™ View Sequencing kits (ThermoFisher Scientific) and 4 barcoded libraries were combined per Ion 316™ Chip kit (ThermoFisher Scientific). Data were analysed using MG-RAST and the number of reads per phylum, class, order, family or genera of species of interest were normalized against all bacteria present. Sequencing runs can be accessed using MG-RAST IDs mgl675297, mgl675300, mgl675312, mgl675291, mgl675309, mgl675318, mgl675306, mgl675294, mgl675321, mgl675315, mgl675285, mgl675288, mgl675324, mgl675303.

### Machine Learning

To identify the metabolic and inflammatory variables most robustly associated with ES-62 treatment, sex, and diet, each of those groups was treated as classes in supervised machine learning classification. Predictors, or input matrix, consisted of 113 pathophysiological, immunological, and metabolic features determined from our cross-sectional chow- and HCD-fed male and female cohorts (comprising 158 mice, 79 males and 79 females). To preserve sample sizes, missing values were imputed using the median of all the available values from other mice for that feature. Each feature was then standardised by centring it around the mean and scaling to unit variance. Models were trained using 5-fold cross-validation on a 75% split of the full dataset, and tested on the remaining unseen 25%. The performance of seven common algorithms was compared (logistic regression, naive Bayes, support vector machines, k nearest neighbours, decision tree, random forests, and gradient boosted trees). We chose to further tune the gradient boosted trees models for inference. Tuning of the gradient boosted trees entailed five rounds of repeated stratified 5-fold splits of the training set in each of which there was a randomised search of parameter space for column sampling size, learning rate, minimum child weight, number of estimators, and maximum tree depth. This generated an ensemble of tuned models from which we selected the best 15 models based on their accuracy in correctly classifying mice from the test set. We then used the resulting distribution of model outputs for ranking the importance of features (Gini index) in correctly classifying the test set mice. Machine learning was performed in Scikit-Learn^76^ and associated plots using Seaborn^77^.

### Statistical Analysis

All data were analysed using GraphPad Prism 6 or 8 software using unpaired student T-tests, one or two-way ANOVA with Fishers LSD post-test for parametric data or Kruskal-Wallis test and Dunn’s post-test for non-parametric data. For clarity, only significant differences between the HCD-PBS and HCD-ES-62 cohorts are shown on the figures, where significance is denoted by *p < 0.05, **p < 0.01 and ***p < 0.001.

### Data Availability Statement

The data supporting the findings are all contained within the article and Supplementary Information files. In addition, the metagenomic data are available from the MG-RAST database using the accession numbers provided in the Methods section. Other primary data files are available from the corresponding authors on reasonable request.

## Acknowledgements

The study was funded by linked awards to MMH and CS (BB/M029727/1) and WH (BB/M029662/1) from the Biotechnology and Biological Sciences Research Council.

## Author contributions

JC, FL, JD, MB, AT, MAP, CL and LM performed the experiments for the study designed by MMH, WH, CS and PAH. JD and FL manufactured ES-62. SB performed the Machine Learning and associated statistical analysis. MMH, WH and CS wrote the paper and all authors were involved in reviewing and revising the manuscript.

## SUPPLEMENTARY INFORMATION

**Supplementary Figure 1:**
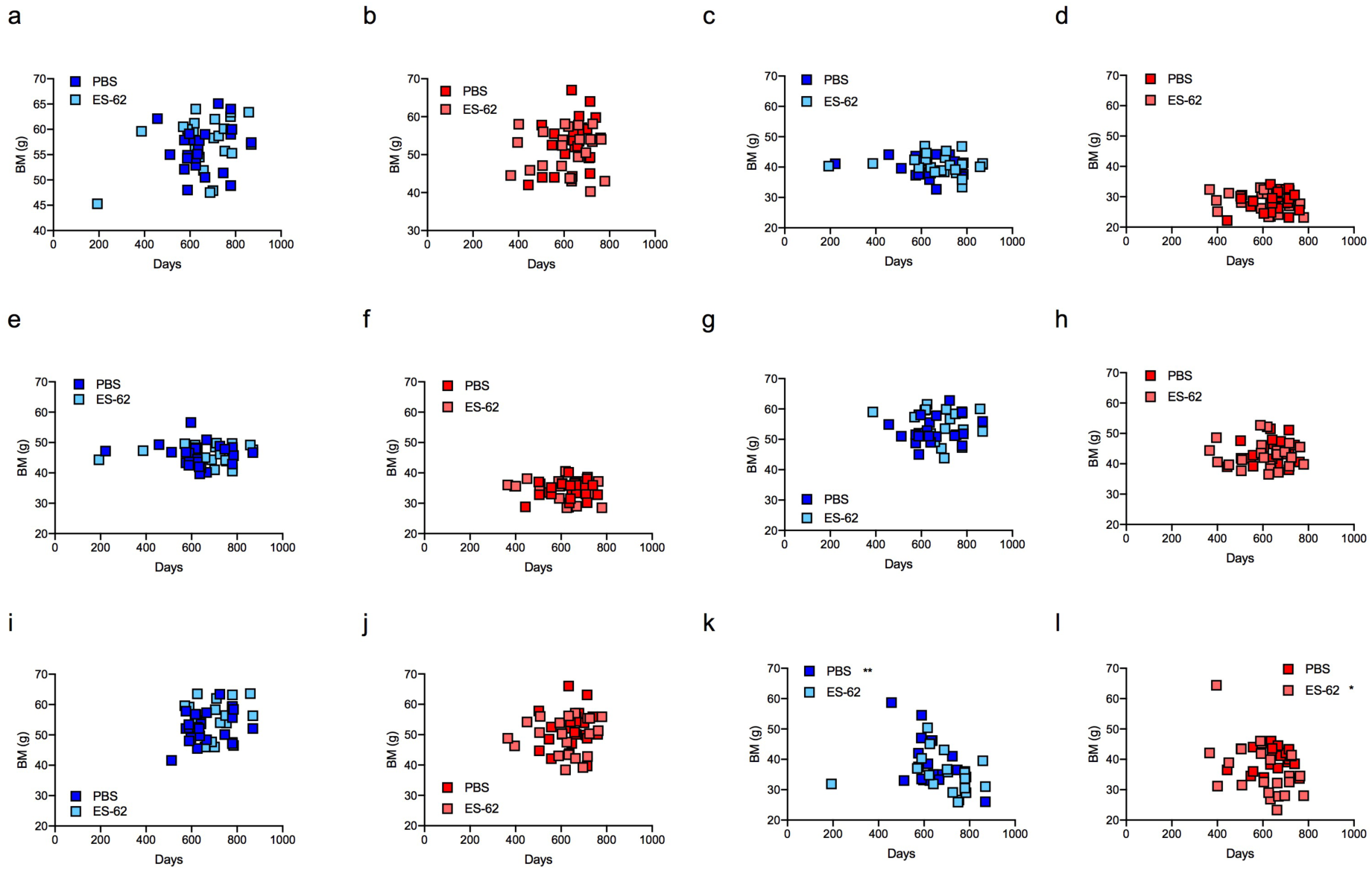
Correlating body mass with age. Correlation of age of death of male (a, c, e, g, i, k) and female (b, d, f, h, j, l) HCD- (PBS- and ES-62-treated) mice with (a, b) peak BM achieved; (c, d) BM at d116; (e, f) BM at d160; (g, h) BM at d340; (I, j) BM at d500; (k, l) BM at death (Male HCD-PBS, r=-0.6053, **p<0.01; Female HCD-ES-62, r=-0.4632, *p<0.05).

**Supplementary Figure 2:**
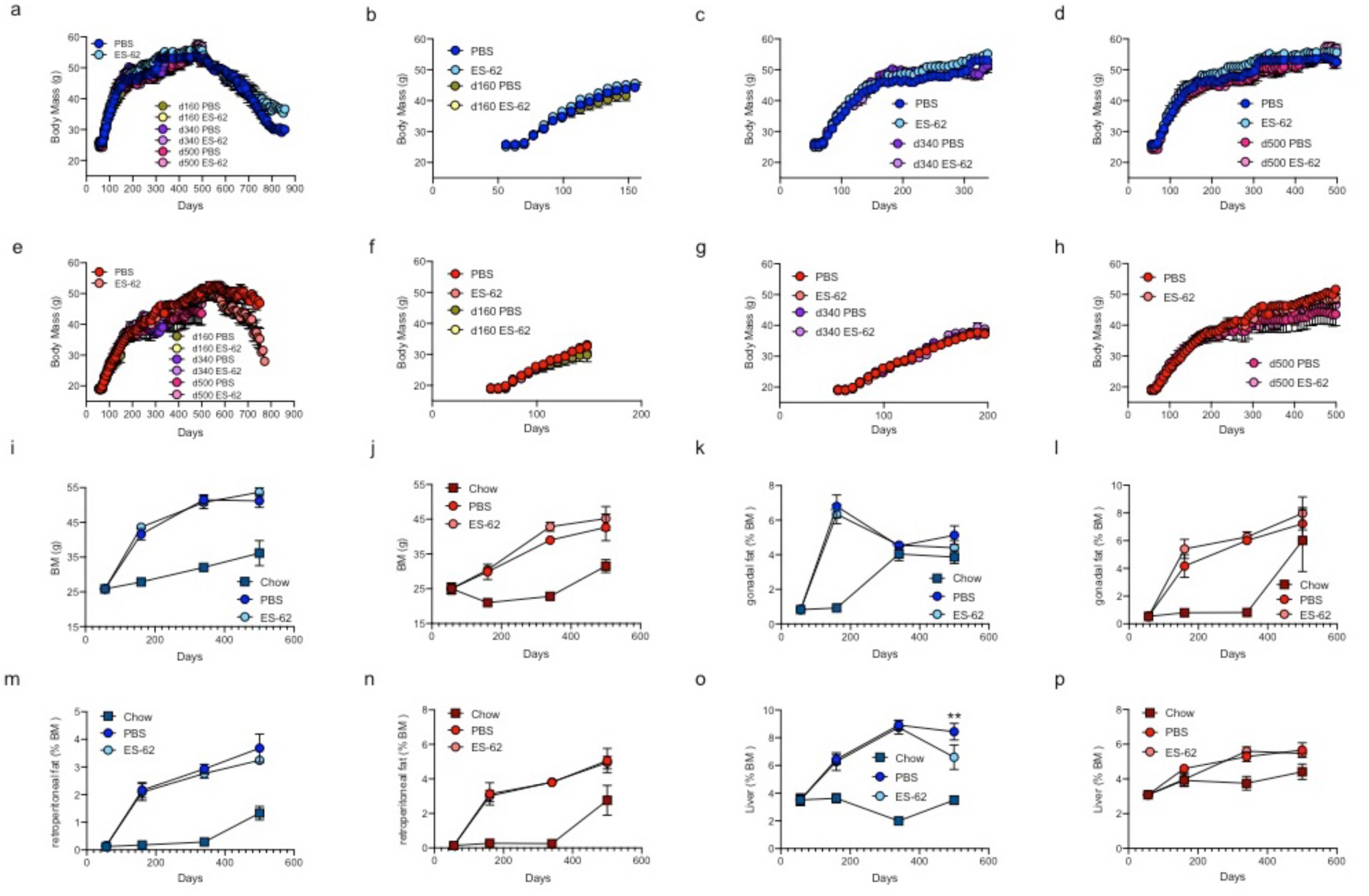
Body mass of survival and cross-sectional cohorts. Longitudinal analysis of BM measurements of the individual male (a-d) and female (e-h) HCD-fed mice in the survival and indicated cross-sectional cohorts where data are presented as mean values ± SEM of the mice surviving or culled at the indicated timepoints. HCD-male and -female mice exhibited substantially higher Body Mass (BM) than their aged-matched chow control groups and male mice were significantly heavier at cull than their female counterparts (***p<0.001) at each timepoint (i & j). Analysis of gonadal fat (% BM; k, l), retroperitoneal fat (% BM; m, n) and liver (% BM; o, p) in male (i, k, m, o) and female (j, l, n, p) cross-sectional cohorts of chow (d56, n=6; d160, n=6; d340, n=6 and d500, n=5) and PBS- or ES-62-treated HCD mice (d160, n=10; d340, n=12 and d500, n=6) at cull. Data are presented as mean values ± SEM of individual mice.

**Supplementary Figure 3:**
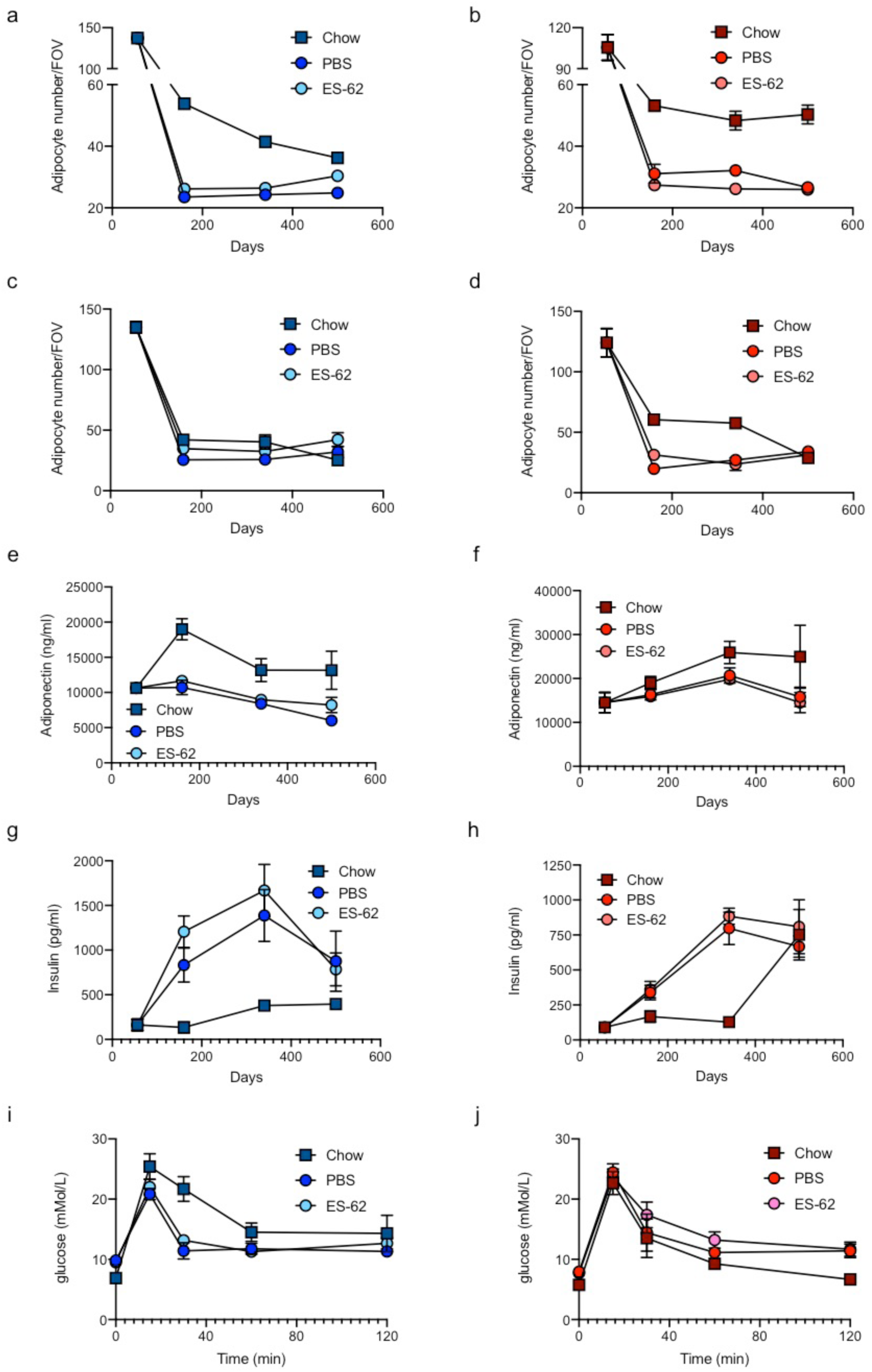
HCD modulation of adipocyte function. Hyperplasia of (small) adipocytes was not detected, as evidenced by quantitative analysis of adipocyte number/FOV in representative images of gonadal (a, b) and retroperitoneal (c, d) adipose tissue sections where data are presented as the mean values ± SEM where n=4-5 individual male (a, c) and female (b, d) chow- and HCD- (PBS- and ES-62-treated) mice at each time point and the values for each mouse are means derived from n=3 replicate analyses. Serum levels of adiponectin (e, f) and insulin (g, h) from male (e, g) and female (f, h) chow- or HCD- (PBS- or ES-62-treated) mice at cull are shown at the indicated time-points. Data are expressed as mean values ± SEM of individual mice. Male cohort sizes: chow - d56, n=6; d160, n=6; d340, n=6; d500, n=5 (adiponectin), 4 (insulin); HCD-PBS - d160, n=9; d340, n=9; d500, n=6; HCD-ES-62 - d160, n=9; d340, n=9; d500, n=6 and female cohort sizes: chow - d56, n=6; d160, n=6; d340, n=6; d500, n=5; HCD-PBS - d160, n=9; d340, n=9; d500, n=6; HCD-ES-62 - d160, n=9; d340, n=9; d500, n=6. Glucose tolerance tests (GTT) were undertaken on male (i) and female (j) chow- and HCD- (PBS- or ES-62-treated) mice one week prior to all culls, with the time course for glucose clearance for the d500 cohorts presented. Data are presented as the mean blood glucose levels ± SEM of individual mice where male (i) cohort sizes: chow - n=5; HCD-PBS - n=6; HCD-ES-62 - n=6 and female (j) cohort sizes: chow - n=3; HCD-PBS - n=6; HCD-ES-62 - n=6.

**Supplementary Figure 4:**
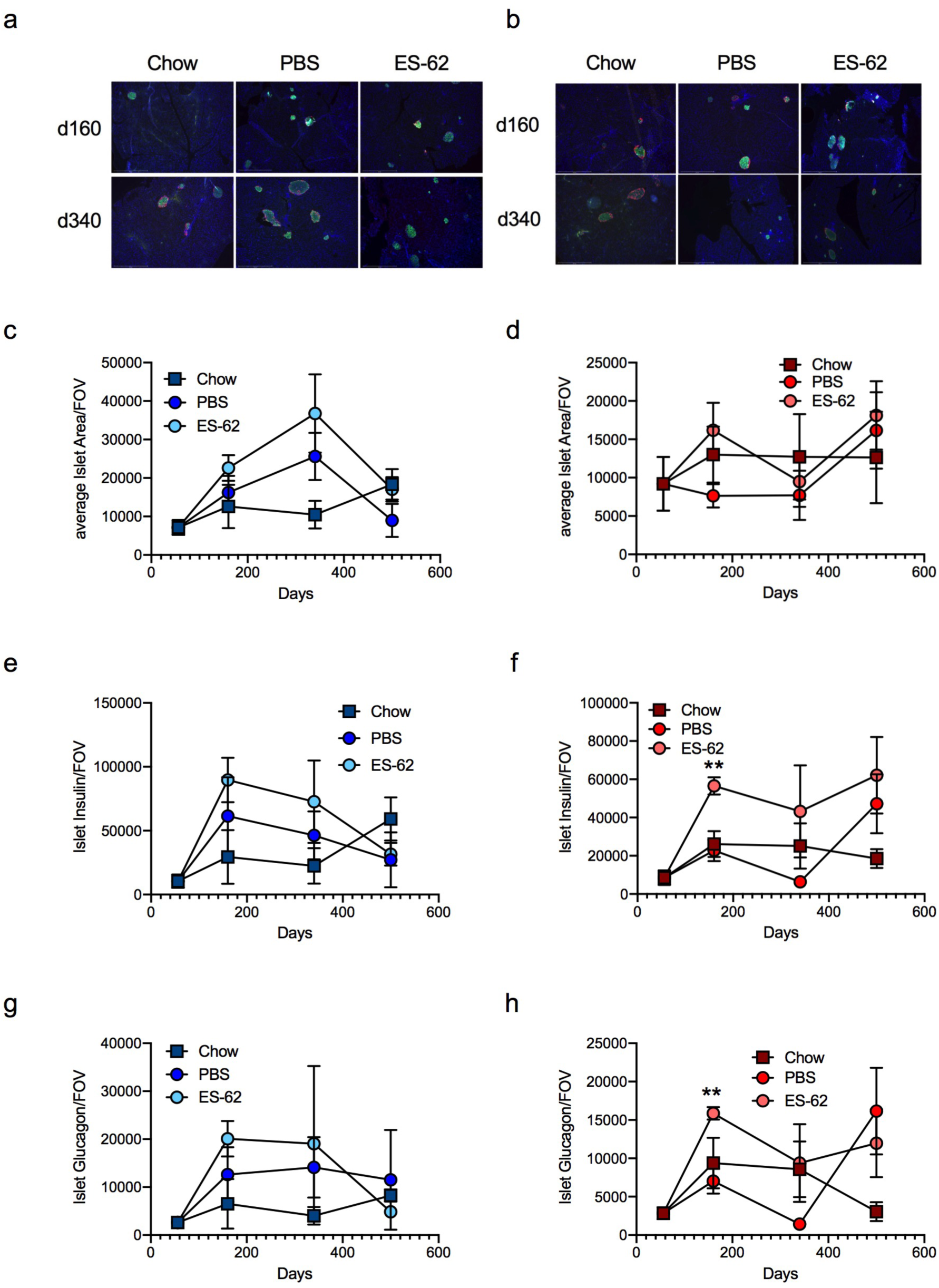
HCD modulation of pancreatic function. Representative images (scale bar 500 µm) of pancreas at d160 and d340 from male (a) and female (b) mice stained for insulin (green), glucagon (red) and counterstained with DAPI (blue). Quantitative analysis of islet size (c, d) and production of insulin (e, f) and glucagon (g, h) where data are presented as the mean values (of triplicate analyses) ± SEM of individual male (c, e, g) and female (d, f, h) mice (n=4-6) at each time-point. For clarity, only significant differences between the HCD-PBS and HCD-ES-62 cohorts are shown on the figures, where significance is denoted by *p < 0.05, **p < 0.01 and ***p < 0.001.

**Supplementary Figure 5:**
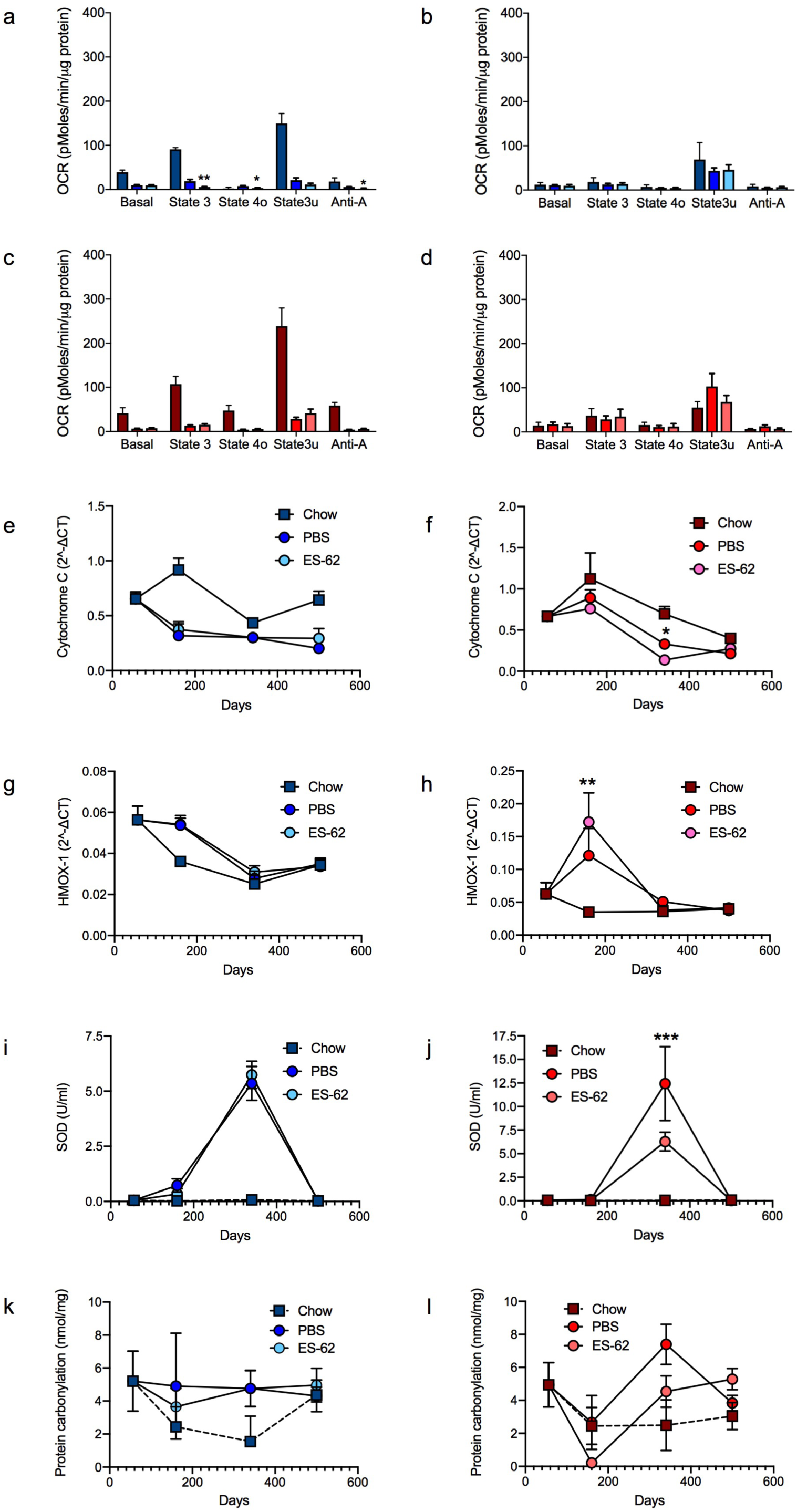
Dysfunctional liver function in HCD-aged mice. Mitochondrial respiration (oxygen consumption rate, OCR) was measured in livers from male (a, b) and female (c, d) chow- or HCD (PBS or ES-62-treated) mice at cull at d340 (a, c) and d500 (b, d). OCR was measured under Basal (substrate alone), State 3 (ADP), State 4 (oligomycin), State 3u (FCCP) and non-mitochondrial (antimycin A plus rotenone; Anti-A) conditions. Data are presented as the values ± SEM of individual mice where cohort sizes were: male, d340 - chow n=6, HCD-PBS n=7, HCD-ES-62, n=8; d500 chow - n=4, HCD-PBS, n=4; HCD-ES-62, n=4; female, d340 - chow n=6; HCD-PBS, n=11; HCD-ES-62, n=12; d500 – chow n=3, HCD-PBS n=4, HCD-ES-62, n=4. RT-qPCR analysis of cytochrome C (e, f) and HMOX-1 (g, h) expression in liver from male (e, g) and female (f, h) chow- and HCD- (PBS- or ES- 62-treated) mice are shown where data are expressed as mean 2^ΔCT values ± SEM of individual mice and the values for each mouse are means of n=3 replicate analyses. Male cohort sizes: chow - d56, n=6; d160, n=6; d340, n=6; d500, n=5; HCD-PBS - d160, n=10; d340, n=11; d500, n=6; HCD-ES-62 - d160, n=10; d340, n=12; d500, n=6 and female cohort sizes: chow - d56, n=6; d160, n=6; d340, n=6; d500, n=5; HCD-PBS - d160, n=5; d340, n=11; d500, n=6; HCD-ES-62 - d160, n=5; d340, n=12; d500, n=6. Determination of superoxide dismutase (SOD; i, j) and protein carbonylation (k, l) activities in liver from male (i, k) and female (j, l) chow- and HCD- (PBS- or ES-62-treated) mice where data are expressed as mean activity values ± SEM of individual mice. Male cohort sizes: chow - d56, n=6 (SOD), n=3 (protein carbonylation); d160, n=6 (SOD), n=3 (protein carbonylation); d340, n=6 (SOD), n=3 (protein carbonylation); d500, n=5; HCD-PBS - d160, n=10 (SOD), n=6 (protein carbonylation); d340, n=10 (SOD), n=11 (protein carbonylation); d500, n=6; HCD-ES-62 - d160, n=8 (SOD), n=6 (protein carbonylation); d340, n=10 (SOD), n=12 (protein carbonylation); d500, n=6 and female cohort sizes: chow - d56, n=5 (SOD), n=3 (protein carbonylation); d160, n=6 (SOD), n=3 (protein carbonylation); d340, n=6 (SOD), 3 (protein carbonylation); d500, n=4; HCD-PBS - d160, n=9 (SOD), 6 (protein carbonylation); d340, n=10 (SOD), n=9 (protein carbonylation); d500, n=6; HCD-ES-62 - d160, n=10 (SOD), n=6 (protein carbonylation); d340, n=10 (SOD), n=11 (protein carbonylation); d500, n=6. For clarity, only significant differences between the HCD-PBS and HCD-ES-62 cohorts are shown on the figures, where significance is denoted by *p < 0.05, **p < 0.01 and ***p < 0.001.

**Supplementary Figure 6.**
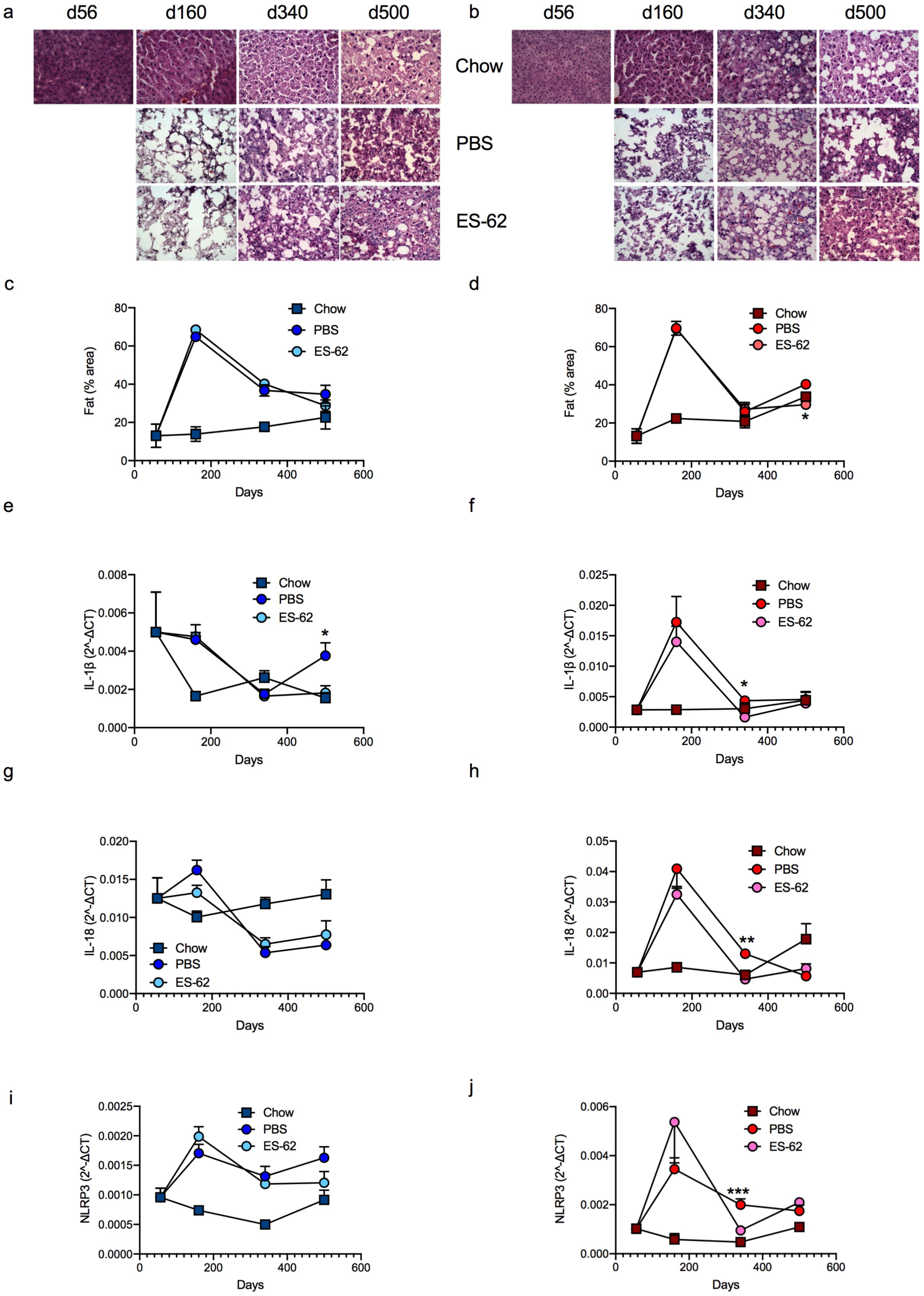
HCD-fed mice exhibit liver steatosis. Representative images (scale bar 100 µm) of liver from male (a) and female (b) chow- and HCD- (PBS- or ES-62-treated) mice stained with H & E and resultant quantitative analysis of fat deposition where data are presented as the mean values ± SEM where n=5-6 individual male (c) and female (d) mice at each time point and the values for each mouse are means derived from n=3 replicate analyses. RT-qPCR analysis of IL-1β, IL-18 and NLRP3 expression in liver from male (e, g, i) and female (f, h, j) chow- and HCD- (PBS- or ES-62-treated) mice where data are expressed as mean 2^ΔCT values ± SEM of individual mice and the values for each mouse are means of n=3 replicate analyses. Male cohort sizes: chow - d56, n=6; d160, n=6; d340, n=6; d500, n=5; HCD-PBS - d160, n=10; d340, n=11; d500, n=6; HCD-ES-62 - d160, n=10; d340, n=12; d500, n=6 and female cohort sizes: chow - d56, n=6; d160, n=6; d340, n=6; d500, n=5; HCD-PBS - d160, n=5; d340, n=11; d500, n=6; HCD-ES-62 - d160, n=5; d340, n=12; d500, n=6.

**Supplementary Figure 7:**
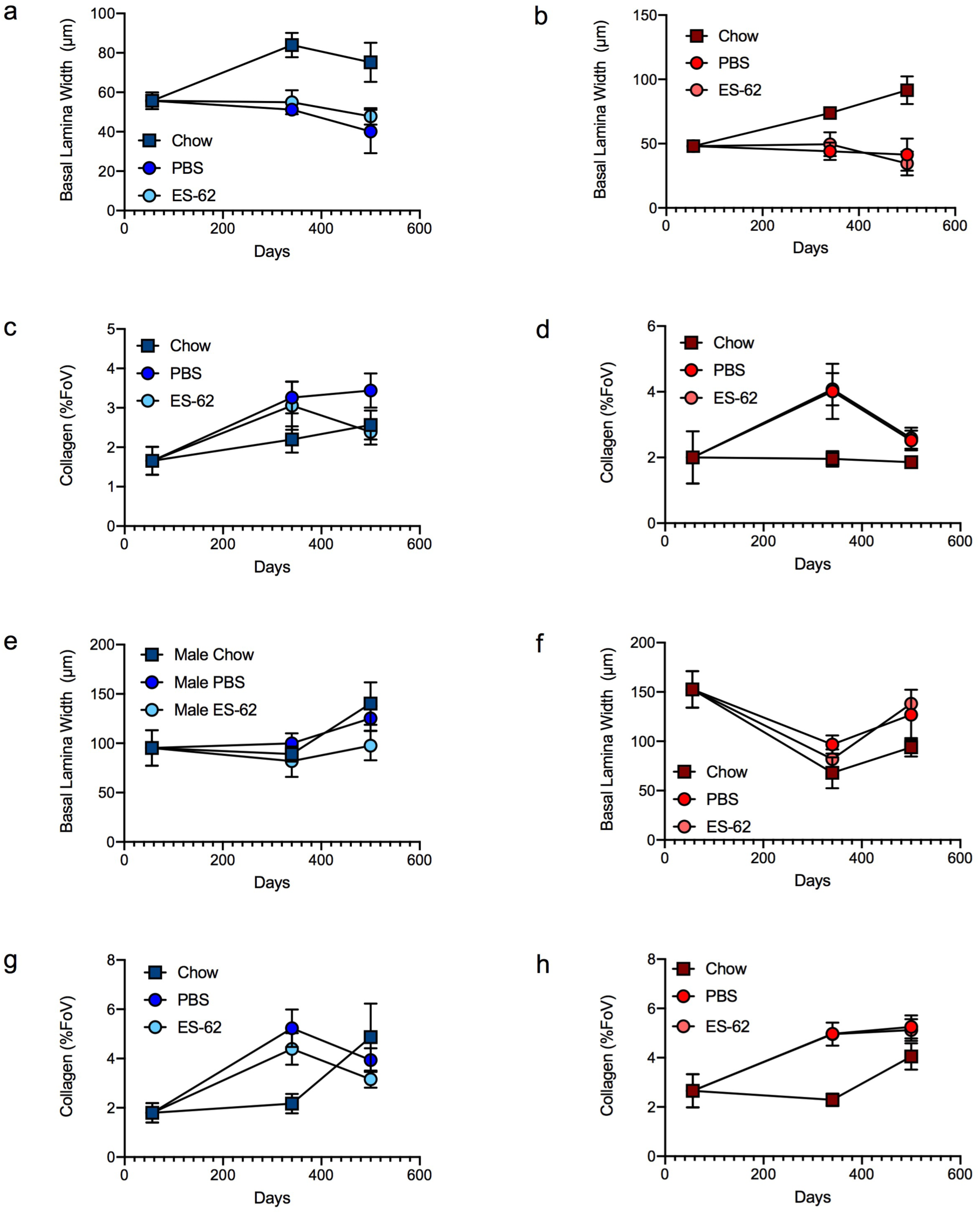
HCD accelerates ageing associated gut damage. Images of ileum and colon tissue from male and female chow- and HCD- (PBS- or ES-62-treated) mice stained with H & E (to visualise basal lamina) and Gömöri’s Trichrome (to visualise collagen deposition) analysed quantitatively for basal lamina width (ileum, a, b; colon, e, f) and collagen deposition (ileum, c, d; colon, g, h) are shown. Data are presented as the mean values ± SEM where n=4-6 individual male (a, c, e, g) and female (b, d, f, h) mice from each group at each time point and the values for each mouse are means derived from n=3 replicate analyses.

**Supplementary Figure 8:**
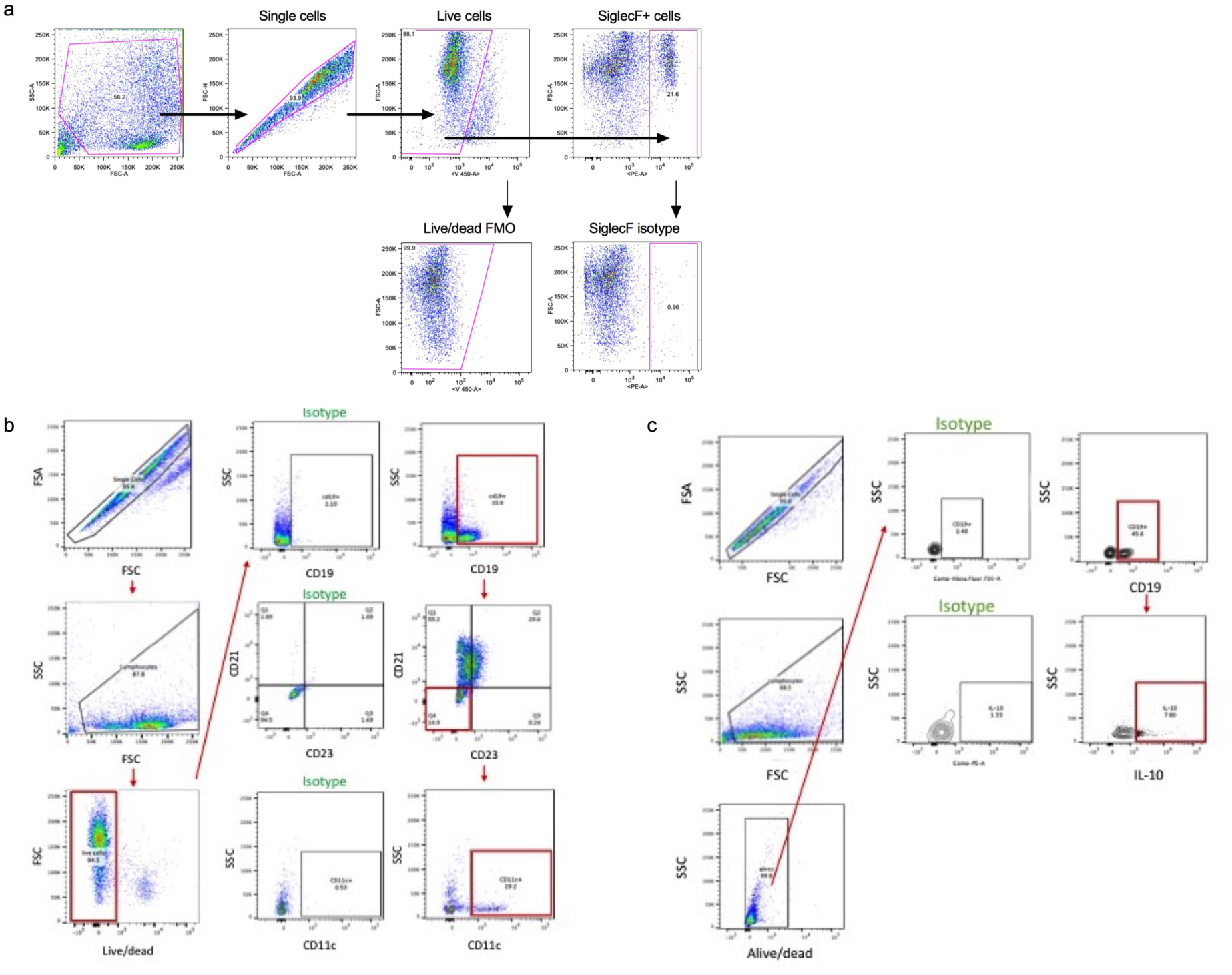
Gating strategies. (a) Following exclusion of dead cells/cell debris and gating of fat cells (by forward scatter versus side scatter), cell doublets were excluded prior to subsequent gating of SiglecF+ populations relative to the relevant isotype and FMO controls (Figure 3). Splenocytes were gated for singlets (FSC-H vs. FSC-A), morphology (FSC-A vs. SSC-A) and then live cells determined by their uptake of the fixable live/dead cell stain before gating prior to assessing expression of (b) CD19^+^CD21^-^CD23^-^CD11c^+^ B cells (Figure 6) or (c) CD19^+^ IL-10^+^ B cells (Figure 6) with reference to relevant isotype controls.

**Supplementary Table 1:** Descriptive Statistics of the longevity cohort

| **Cox Regression Analysis<sup>a</sup>** |  |  |  |  |  |  |
| --- | --- | --- | --- | --- | --- | --- |
| **Covariate** | **B** | **SE** | **Wald** | **Df** | **Sig** | **Exp(B)** |
| Treatment | -0.130 | 0.206 | 0.399 | 1 | 0.528 | 0.878 |
| Sex | 0.541 | 0.222 | 5.933 | 1 | 0.015 | 1.717 |
| **Descriptive Statistics<sup>b</sup>** |  |  |  |  |  |  |
| **Sex** | **Treatment** | **Median** | **Mean** | **Min-Max** | **Oldest 10%** | **n** |
| Male | PBS | 629 | 636 ± 26.7 | 222-869 | 782 | 24 |
|  | ES-62 | 703 | 676 ± 29.8 | 193-869 | 821 | 24 |
| Female | PBS | 653 | 637 ± 16.7 | 443 -760 | 727 | 24 |
|  | ES-62 | 629 | 610 ± 23.9 | 366 -779 | 745 | 24 |

<sup>a</sup>Cox regression analysis of PBS- and ES-62 -treated mice. Age at death was analysed in pooled male and female PBS and ES-62 mice using Cox regression analysis (SPSS Statistics v22). The independent variables, Treatment (PBS or ES-62) and Sex (male or female), were replaced with a set of indicator variables to denote the presence or absence of category membership. B is the un-standardised regression coefficient and its standard error is SE, its Wald test statistic value, Wald, the degrees of freedom, df and the significance, Sig. Exp(B) for the covariate of interest is the predicted change in the hazard ratio for a unit increase in the predictor. Sample size = 48 for PBS- and 48 for ES-62-treated mice. <sup>b</sup>Descriptive statistics of this longevity study were derived using the GraphPad Prism8 logrank test survival software package.

**Supplementary Table 2:**
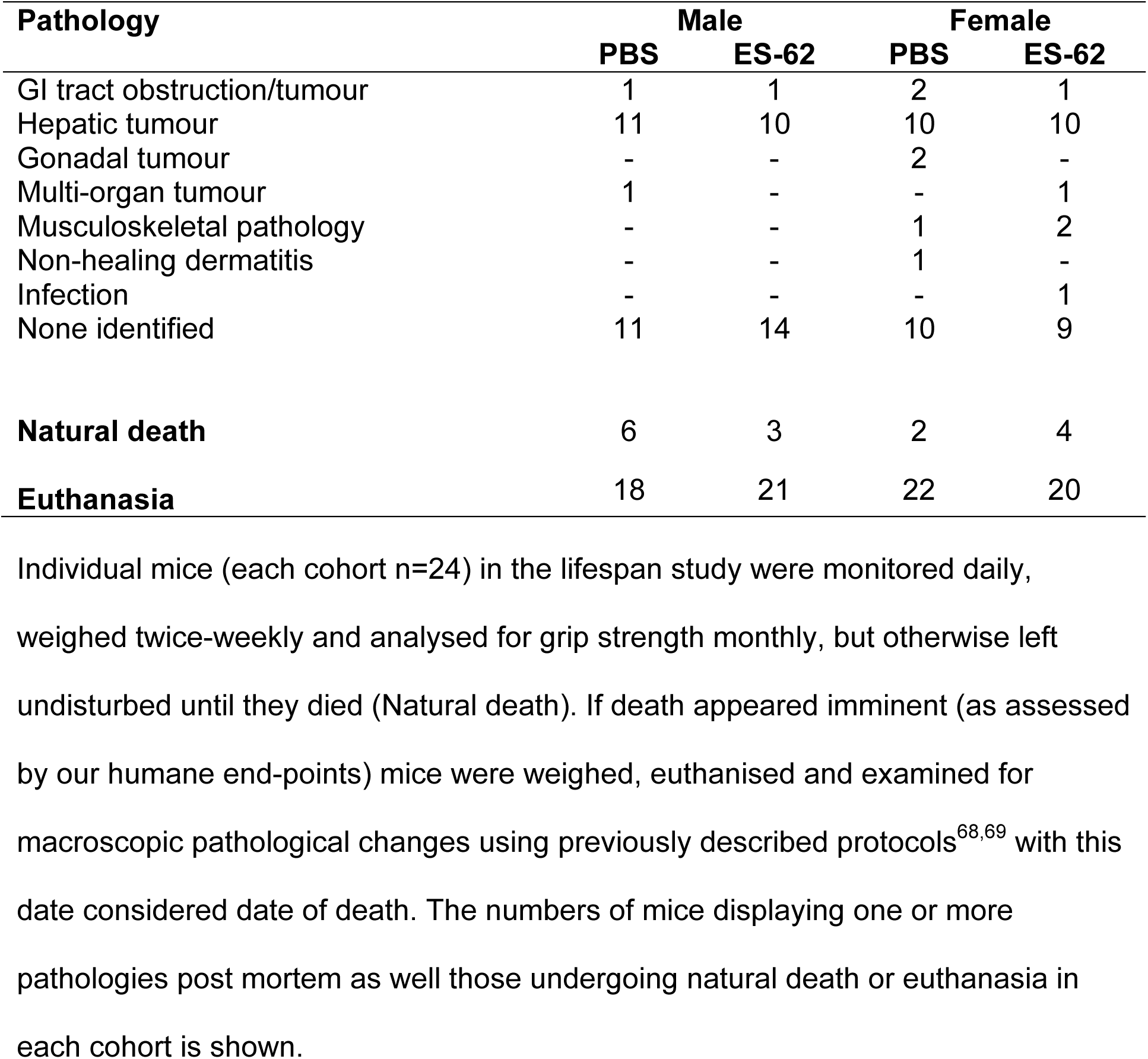
Pathology identified post-mortem in mice from the lifespan cohort

